# Histone lactylation: a new epigenetic mark in the malaria parasite *Plasmodium*

**DOI:** 10.1101/2024.08.19.608576

**Authors:** Ibtissam Jabre, Nana Efua Andoh, Juliana Naldoni, William Gregory, Haddijatou Mbye, Chae Eun Yoon, Aubrey J. Cunnington, Athina Georgiadou, Andrew M. Blagborough, Catherine J. Merrick

## Abstract

Epigenetic processes play important roles in the biology of the malaria parasite *Plasmodium falciparum*. Here, we characterised a new epigenetic mark, histone lactylation, recently discovered in humans: it was found in two human malaria parasites, *P. falciparum* and *P. knowlesi*, and also *in vivo* in two rodent malaria models. Histones were lactylated rapidly in response to elevated lactate levels, and rapidly delactylated when lactate levels fell. Thus, this mark is well-placed to act as a metabolic sensor, since severe falciparum malaria characteristically leads to hyperlactataemia. Mass spectrometry showed that lysines on several parasite histones could be lactylated, as well as many non-histone chromatin proteins. Histone lactylation was less abundant and less inducible in *P. knowlesi* than *P. falciparum*, suggesting that *P. falciparum* may have evolved particular epigenetic responses to this characteristic feature of its pathology. Finally, in the rodent model *P. yoelii*, hyperlactataemia correlated with parasite transcriptomic programmes that suggested metabolic ‘dormancy’.

## INTRODUCTION

Human malaria occurs when protozoan *Plasmodium* parasites infect red blood cells. Six *Plasmodium* species can cause malaria in humans, including the main agent of severe and fatal malaria, *P. falciparum*, and the zoonotic macaque parasite *P. knowlesi.* These two are currently the only human malaria parasites that can be cultured continuously *in vitro*.

Clinical malaria has many characteristic pathological features, one of which is hyperlactataemia. This is clinically defined as a level of blood lactate above 5mM, but patients with falciparum malaria can reach at least 15mM ^1,2^. Parasites in the bloodstream respire by glycolysis, producing lactate, and they also cause their host erythrocytes to adhere to capillary walls, impeding blood flow and promoting anaerobic respiration in host tissues. Collectively, this results in hyperlactataemia, which is predictive of severe disease and death, as it can lead to the fatal malaria syndrome of metabolic acidosis and respiratory distress ^3^.

*Plasmodium* parasites could benefit from being able to sense the metabolic environment in their host and modify virulence accordingly – for example, by increasing commitment to sexual stages that are competent for transmission to new hosts, or by modulating cytoadherence of infected erythrocytes. This fascinating area of host-parasite biology is increasingly being explored, with aspects of blood chemistry including levels of glucose, magnesium and S-adenosylmethionine being proposed to influence the virulence of *P. falciparum* parasites ^4–7^.

Blood lactate has yet to be fully explored in this context, but since it correlates with parasite load and disease severity, it could potentially serve as both a ‘quorum sensor’ and a measure of host stress. In one human infection study, lactate was linked to the expression pattern of a key virulence gene family, *var*, which encodes variant adhesins expressed on infected erythrocytes ^1^. The *var* gene family is known to be epigenetically controlled by histone acetylation and methylation. Epigenetics also plays other prominent roles in *Plasmodium* biology, controlling aspects of virulence including alternate invasion pathways and sexual stage commitment (gametocytogenesis) ^8^. In a recent *in vitro* study, gametocytogenesis was also linked to lactate, with parasites cultured in added lactate converting to gametocytes at an elevated rate ^9^.

Here, we explored lactate as a potential epigenetic modifier in *Plasmodium*, following the recent discovery of lactyl-lysine as a new epigenetic mark in mammalian cells, similar to the well-characterised acetyl-lysine ^10^. Since its discovery in 2019, this epigenetic mark has been heavily explored in the context of human cells in high-lactate environments – primarily tumours experiencing the Warburg effect ^11,12^. It has not yet been explored in the context of malaria, a disease characterised by systemic hyperlactataemia; however, we did recently observe that histone lactylation could occur in *P. falciparum*, and hence proposed this as a new virulence modifier in *Plasmodium* ^13^. We now report that the epigenetic mark is conserved in several *Plasmodium* species, occurring both *in vitro* and *in vivo*. It appears on many histone lysine residues, is rapidly labile and is responsive to lactate levels both in culture and in mammalian host blood. Therefore, it could have many virulence-modulating roles.

## RESULTS

### *Plasmodium* histones are inducibly lactylated

We first measured the level of lactyl-lysine in *Plasmodium* histones after treating cultured parasites with increasing levels of lactate. Trophozoite-stage *P. falciparum* parasites were exposed to 5-25mM lactate, added to the culture media for 12h. **Figure 1A** shows that their histones were inducibly and titratably lactylated, with histone H4 showing the strongest lactyl-lysine signal. An antibody identifying a single lactylated residue on histone H4 (K12) confirmed that the general lactyl-lysine antibody recognised this histone, but lactylation of the K12 residue was not strongly inducible **(fig 1B)**. We conducted the same experiments on a second species, *P. knowlesi*, and showed that histone lactylation was similarly inducible **(fig 1C)**. It was, however, much less abundant than in *P. falciparum*, since greater amounts of parasite extract were required to detect similar signals on *P. knowlesi* histones, and the specific residue H4K12La was barely detectable **(fig 1D)**.

**1.**
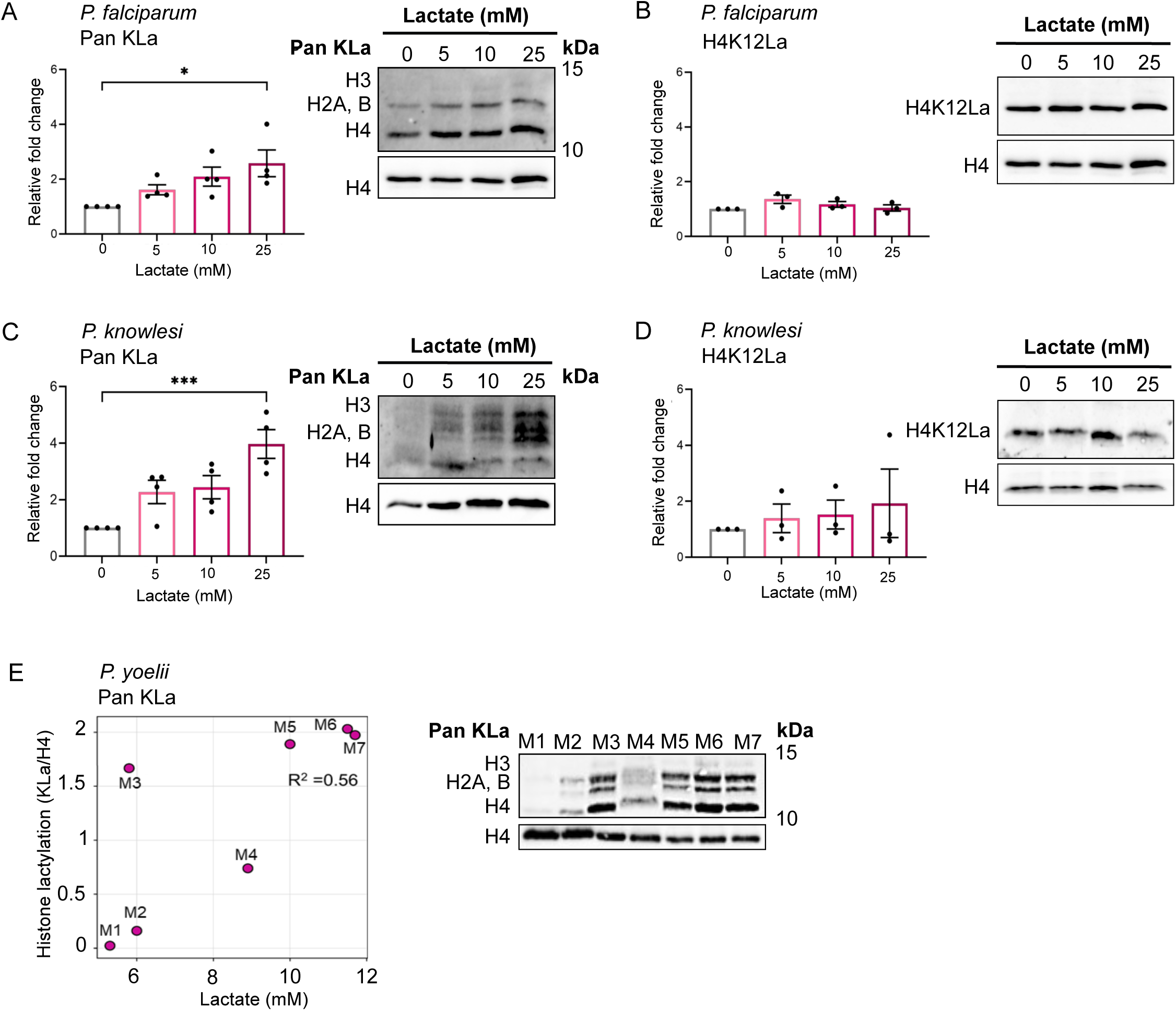
*Plasmodium* histones are inducibly lactylated. A, B: Lysine lactylation was measured in *P. falciparum* parasites grown in different concentrations of L-lactate for 12h. Western blots show ‘Pan KLa’(A) and histone H4 K12 lactylation (‘H4K12La’, (B)), as well as total histone H4 as a control. Graphs show quantifications of blots from several independent experiments (n=4 for PanKLa, n=3 for H4K12La) with each lactyl-histone signal quantified relative to histone H4, then expressed as a fold-change from the 0mM condition (bars show means, dots show individual values, error bars show SEM). Multiple comparison of means was performed using one-way ANOVA and Tukey HSD post hoc test: *, p < 0.05; ***, p < 0.001; comparisons not shown, not significant. A representative western blot from each group of experiments is shown. C, D: Lysine lactylation was measured in *P. knowlesi* parasites grown in different concentrations of L-lactate for 8h, as in (A, B). E: Lysine lactylation was measured in *P. yoelii* 17XL parasites exposed to varying levels of lactate in the blood of the host mouse. Western blots show ‘Pan KLa’ and histone H4 as control. M1-M7, individual mice. Scatter plot represents Pearson correlation (R^2^= 0.56) between histone KLa signal intensities normalized to H4 (*y-*axis) and lactate levels in the blood (*x-*axis) upon parasite collection from the host.

We then investigated whether this phenomenon occurs naturally *in vivo* during malaria. The rodent model species *P. yoelii*, strain 17XL, can generate severe hyperlactataemia ^14^, and blood lactate levels in mice infected with *P. yoelii* 17XL did indeed correlate with histone lactylation in the parasite **(fig 1E)**. Notably, not all rodent malaria models lead consistently to hyperlactataemia and in the *P. berghei* ANKA strain, which does not ^14^, we did not observe a strong correlation between blood lactate and histone lactylation **(fig S1)**. Overall, these results revealed that there is a conserved pathway for histone lactylation in *Plasmodium,* induced by hyperlactataemia.

### Histone lactylation is rapidly inducible and reversible

We next investigated the time dynamics of the modification: **figure 1** shows that it can occur within 12h and 8h in *P. falciparum* and *P. knowlesi* respectively, but it does not show how fast the modification was actually added, or how fast it would be removed if exogenous lactate levels subsequently fell. To investigate this, parasites were tightly synchronised at rings, washed into fresh media, exposed to a pulse of 25mM lactate when they reached early trophozoites, then followed by western blotting at intervals over the subsequent 20-24h. (25mM lactate, for 12h in *P. falciparum* or 8h in *P knowlesi* due to its shorter cell cycle, was chosen for these and subsequent experiments because, from **figure 1**, this induced the strongest histone lactylation, without markedly affecting parasite growth **(fig S2)**, whereas even higher levels of ≥ 30mM over 48h were previously reported to affect parasite growth in *P. falciparum*^15^).

Figure 2A shows that histone lactylation rose within just 2h of exposure; continued to rise after 6h, and then fell back slightly after 12h. This was surprising, but we surmised that it may be because of a natural cell-cycle-related pattern in histone lactylation, since parasite metabolism, which peaks in trophozoites, produces its own endogenous lactate. This was confirmed by an identical parallel timecourse in which no exogenous lactate was added **(fig 2A**: grey bars). Here, histone lactylation was still induced, but to a much lesser extent, peaking in the mid-trophozoite stage (∼30-32hpi) and falling back slightly in late trophozoites (∼36-38hpi).

**2.**
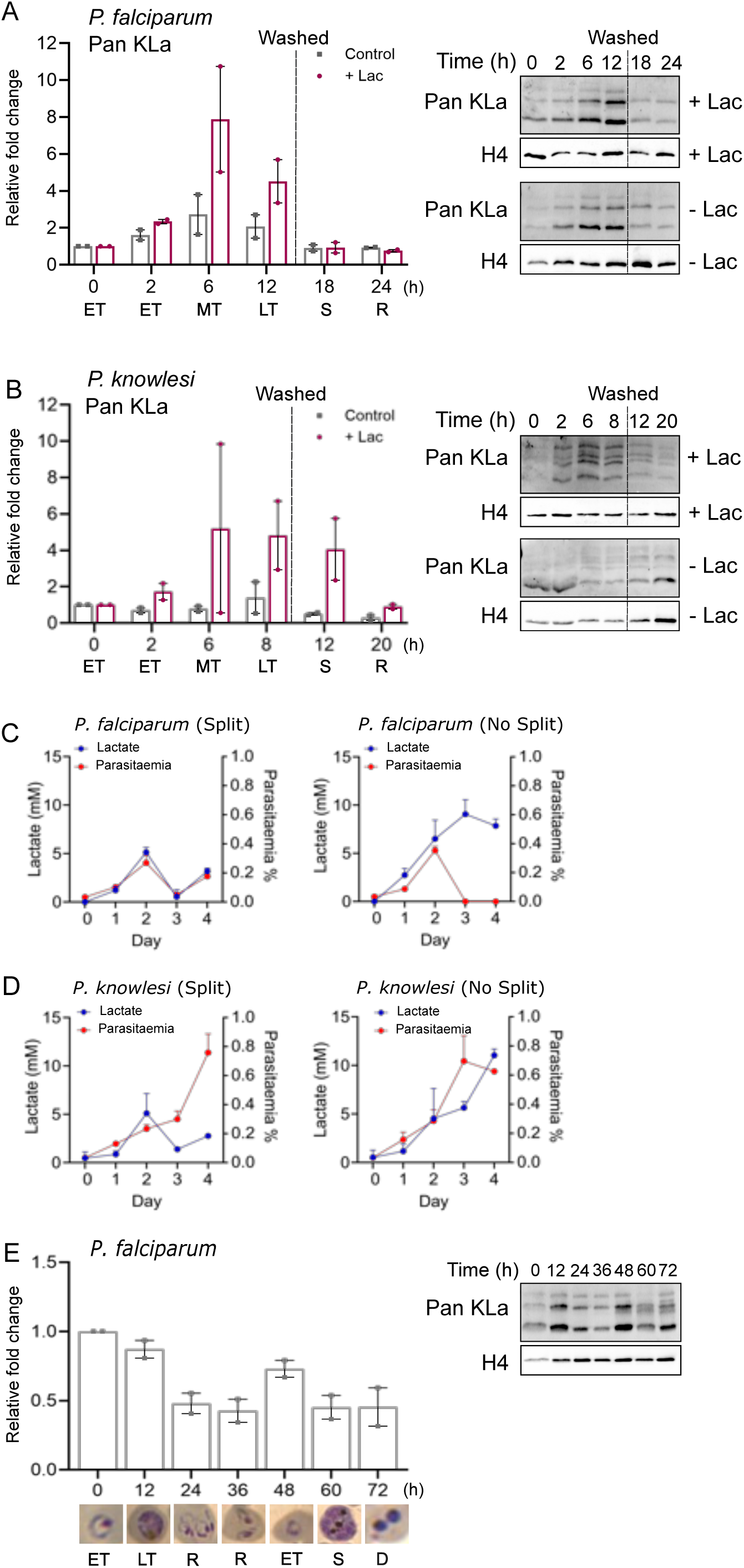
Histone lactylation is rapidly inducible and reversible. A: Lysine lactylation (‘Pan KLa’) measured in *P. falciparum* parasites grown for 12h from the early trophozoite stage either in 25mM L-lactate (+Lac), or in control media without added lactate (Control). Parasites were then washed into fresh media. Samples were taken for western blotting at intervals from 0 to 24h. Stage of the parasite culture is shown on the x-axis: ET, early trophozoite; MT, mid trophozoite; LT, late trophozoite; S, schizont; R, ring. Graph shows data (mean and range) from two independent timecourse experiments, each quantified as fold-changes from time-0 in the signal of KLa versus total-H4, with one representative western blot shown. B: Lysine lactylation (‘Pan KLa’) measured in *P. knowlesi* parasites grown for 8h from the early trophozoite stage either in 25mM L-lactate, or in control media without added lactate, then washed into fresh media. Samples were taken for western blotting at intervals from 0 to 20h. Stage of the parasite culture is shown on the x-axis as in A; data were quantified and represented as in (A). C,D: Lactate levels and parasitaemias were measured every 24h in cultures of *P. falciparum* (C) and *P. knowlesi* (D), starting at 0.5% parasitaemia, either split back to 0.5% every 48h or allowed to grow continuously without splitting. Data shown are the mean of biological duplicate experiments. E: *P. falciparum* parasites were grown from 1% parasitaemia without splitting or changing the media for 72h, starting at the early-trophozoite stage and sampling every 12h. Stage of the parasite culture is shown on the x-axis as in A, and example parasite morphology is also shown. By 60h, morphology showed that the culture was starting to crash, with dead (D) pyknotic parasites by 72h. The graph shows fold-changes in KLa signal from time-0, quantified from biological duplicate experiments, as in (A,B); one representative western blot is shown.

To measure the dynamics of de-lactylation, parasite cultures were washed after 12h and then followed at subsequent timepoints 6 and 12h later (re-washing at each of these points to keep the lactate levels in the media low). By 6h (i.e. in schizonts at ∼42-44hpi), the level of histone lactylation had reverted to baseline levels. Again, this followed the ‘natural’ cell cycle pattern, which was greatly amplified by exposure to exogenous lactate. Thus, histone lactylation was rapidly responsive – in both directions – to changes in the lactate content of the media.

We repeated these experiments in *P. knowlesi* **(fig 2B)**: again, the histone lactylation signal was inducible across the trophozoite stage of the cell cycle, while background levels of lactylated histones were low, variable, and not clearly cell-cycle-regulated in this species **(fig 2B**: grey bars). As in figure 1, the signal was weaker than in *P. falciparum:* 4-5 times the quantity of *P. knowlesi* histones was required to detect comparable levels of histone lactylation by western blotting.

To measure the extent of endogenous lactate production in our parasite culture system, we measured lactate levels in the media, both when parasites were cultured under standard conditions (reducing the parasitaemia and changing the media every 48h), and also when they were allowed to grow continuously to the point of ‘crashing’ (the term for parasite death due to high parasitaemia and exposure to spent media ^16^) **(fig 2C,D)**. As expected, media lactate levels correlated with parasitaemia, reaching only ∼5mM within 2 days as a culture of *P. falciparum* grew from 0.5% to 4.6% parasitaemia **(fig 2C)**. When the culture was allowed to ‘crash’, lactate levels peaked at ∼10mM **(fig 2C)**. The profile of lactate production was similar in *P. knowlesi* cultures, although this parasite is more resistant than *P. falciparum* to ‘crashing’, so these parasites could grow exponentially for at least 3 days, producing lactate continuously and reaching 11.5mM by day-4 **(fig 2D)**. We also followed histone lactylation in *P. falciparum* over 72h by western blotting, confirming that it fluctuated in a stage-dependent manner even as lactate accumulated in the media, with lactylation being consistently higher in trophozoites and lower in rings **(fig 2E).**

Overall, these data show that routine culture of both parasite species can expose them to significant but modest lactate fluctuations. Routine cultures can reach the clinical threshold for hyperlactataemia (5mM), which is sufficient to induce some histone lactylation **(fig 1)**, but is much lower than the levels attained in human patients with severe malaria ^1,2^.

### Histone acetylation is affected by exposure to high lactate

In human cells, both acetylation and lactylation have been detected on the same histone lysine residues, and it has been suggested that histone acetyltransferase enzymes may moonlight as lactylases ^10^. This raised the possibility that in *Plasmodium*, histone lactylation could reciprocally affect histone acetylation – a well-established factor in gene expression regulation.

We tested this by western-blotting for acetyl H4 and acetyl H3 under conditions of high lactate exposure, as in figure 1. Figure 3A-D shows a trend towards lactate-inducible acetylation, with acetylation of H3 rising up to 2-fold after exposure to 25mM lactate and acetylation of H4 rising an average of 2-fold, albeit with less consistency between experiments. Thus, acetylation was less strongly lactate-inducible than lactylation, but the two histone marks could be mutually regulated by lactate.

**3.**
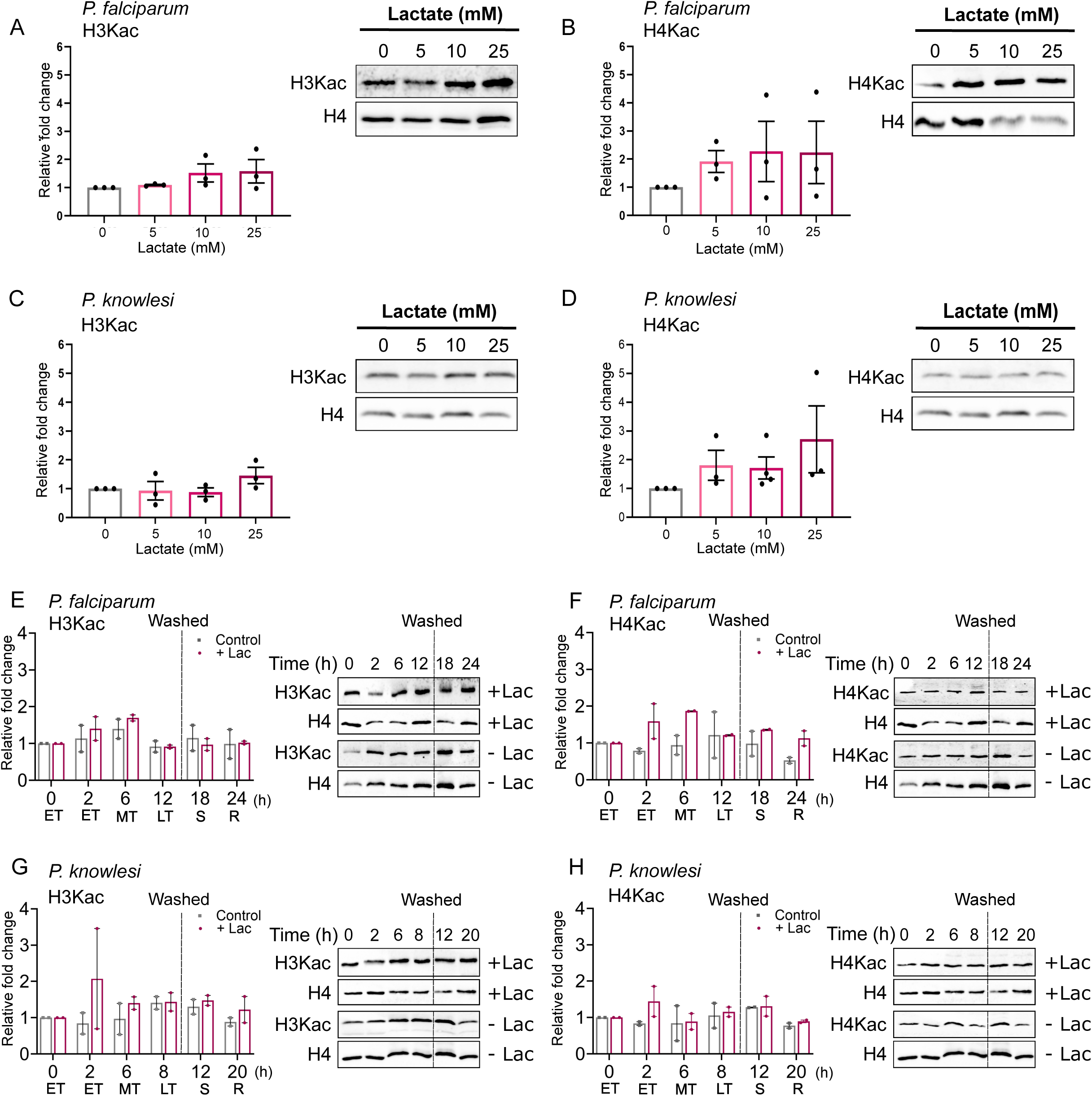
Histone acetylation is affected by exposure to high lactate. A,B: Histone 3 lysine acetylation (‘H3Kac’, (A)) and histone 4 lysine acetylation (‘H4Kac’, (B)) were measured in *P. falciparum* parasites grown in different concentrations of L-lactate for 12h, as in figure 1. Graphs show quantification of blots from several independent experiments (n=3) with each acetyl-histone signal quantified relative to histone H4, then expressed as a fold-change from the 0mM condition (bars show means, dots show individual values, error bars show SEM). Multiple comparison of means was performed using one-way ANOVA and Tukey HSD post hoc test: none were significantly different at p < 0.05. A representative western blot from each group of experiments is shown. C,D: Histone 3 lysine acetylation (‘H3Kac’, (C)) and histone 4 lysine acetylation (‘H4Kac’, (D)) were measured in *P. knowlesi* parasites grown in different concentrations of L-lactate for 8h, as in figure 1; data represented as in (A,B). E,F: Histone 3 lysine acetylation (‘H3Kac’, (E)) and histone 4 lysine acetylation (‘H4Kac’, (F)) were measured in *P. falciparum* parasites grown either in 25mM L-lactate, or in control media without added lactate, for 12h, then washed into fresh media, as in figure 2. Samples were taken for western blotting at intervals from 0 to 24h. Stage of the parasite culture is shown on the x-axis: ET, early trophozoite; MT, mid trophozoite; LT, late trophozoite; S, schizont; R, ring. Graph shows data (mean and range) from two independent timecourse experiments, each quantified as fold-changes from time-0 in the signal of Kac versus total-H4, with one representative western blot shown. G,H: Histone 3 lysine acetylation (‘H3Kac’, (G)) and histone 4 lysine acetylation (‘H4Kac’, (H)) were measured in *P. knowlesi* parasites grown either in 25mM L-lactate, or in control media without added lactate, for 8h, then washed into fresh media, as in figure 2. Samples were taken for western blotting at intervals from 0 to 20h. Stage of the parasite culture is shown on the x-axis; data represented as in (E,F).

We also followed the time dynamics of histone acetylation following lactate exposure, using timecourses as in figure 2 **(fig 3E-H)**. Acetylation did not vary greatly across the normal cell cycle (fluctuations were less than 2-fold) but after exposure to 25mM lactate, histones were, again, additionally acetylated, particular on H4 in *P. falciparum*. This occured as rapidly as lactylation – within 2h – but deacetylation was not equally rapid after lactate removal (this was also seen particularly on H4 in *P. falciparum* **(fig 3F)**). This may suggest that histone acetyltransferase activity is shared for both modifications, whereas deacetylase activity is not.

### Histone lactylation is distributed throughout the nucleus

In *P. falciparum,* certain histone modications are specifically associated with heterochromatin, which tends to be located at the nuclear periphery, marked by heterochromatin protein 1 (HPI) ^17^, whereas other modifications associated with euchromatin appear throughout the nucleoplasm. We investigated the location of histone lactylation using immunofluorescence microscopy **(fig 4)**. We found no evidence for compartmentalisation of lactylated histones, with both the general lactyl-lysine signal **(fig 4A)** and the specific signal for H4K12 **(fig 4B)** appearing throughout the nuclei of all parasite stages. Neither signal colocalised with HP1, and both appeared similar to the pan-nuclear distribution of histone acetylation (**fig S3**). The great majority of lactyl-lysine signal was nuclear **(fig 4A)**, suggesting that there is little lactylated protein in the cytoplasm of *Plasmodium*.

**4.**
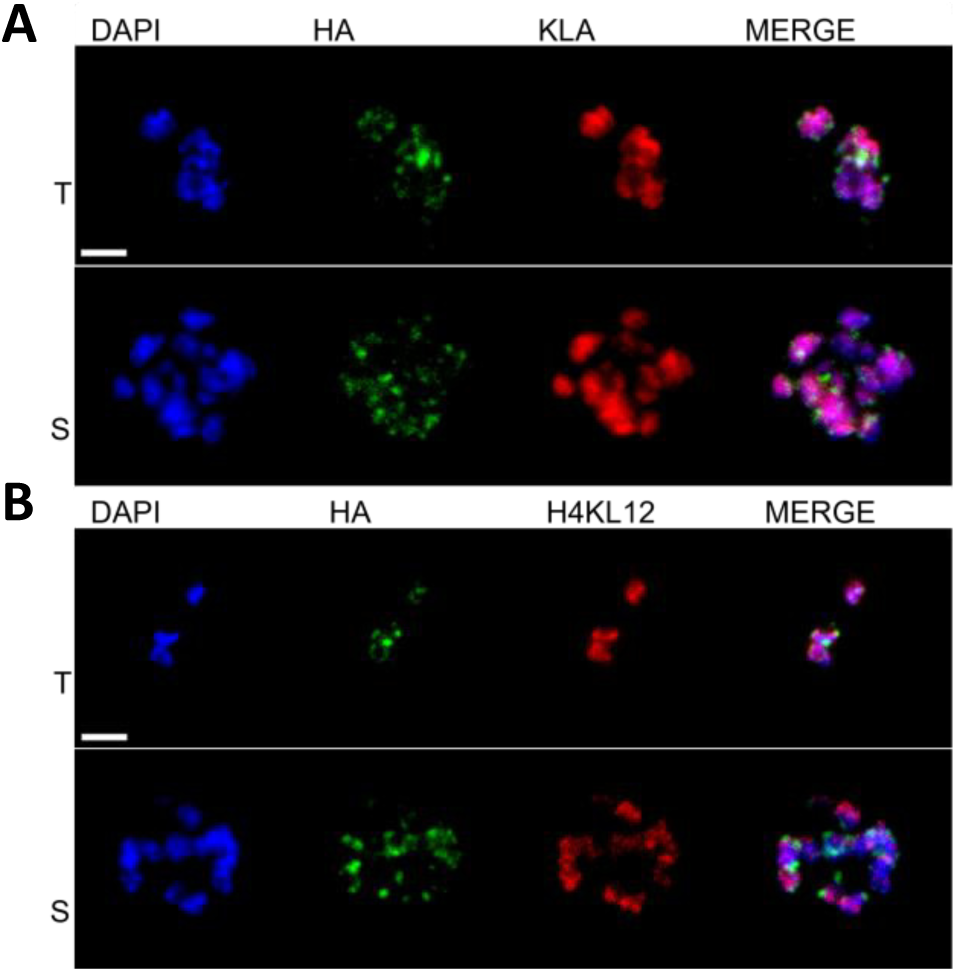
Histone lactylation is distributed throughout the nucleus. Confocal microscopy images showing the location of lactyl PTMs and of HP1 in *P. falciparum* 3D7 HP1_HA. Lactyl-lysine, red; HP1_HA, green; DAPI, blue. KLa (A) and H4K12La (B) distributions are mostly nuclear, and do not colocalise specifically with HA-tagged HP1. T, trophozoite stage; S, schizont stage. Scale bar (2 µm) applies to all images.

### Histone lactylation occurs on multiple histone lysine residues

For a comprehensive view of the histone lysine residues that can be lactylated in both *P. falciparum* and *P. knowlesi*, we used liquid chromatography-tandem mass spectrometry (LC-MS/MS). Trophozoite-stage parasites were either exposed or not exposed to 25mM lactate and their histones were acid-extracted from chromatin in biological triplicate. In a single pilot experiment, histones were also extracted from mixed stage cultures (+/- 25mM lactate), enabling detection of KLa sites in all stages of the asexual cycle. Histone integrity and the induction of lactylation were confirmed via Coomassie-stained SDS-PAGE and by immunoblotting for KLa **(Fig S4)**.

Mass spectrometry on the histone-containing region of the gel identified all eight histones (H2A, H2A.Z, H2B, H2B.Z, H3, H3.3, CenH3 and H4), with sequence coverage >67.5% except in the case of CenH3 **(Tables S1-4).** We then mapped KLa sites, as previously reported ^10^ **(Fig S5A)**, revealing 16 and 21 KLa sites in *P. falciparum* and *P. knowlesi* respectively **(Fig 5A)**. These were unevenly distributed across the histone tails and almost all KLa sites could also be acetylated **(Fig S5B, Tables S1-4).** Interestingly, despite the histone sequences being highly conserved between *Plasmodium* species, the majority of KLa sites were species-specific, suggesting that this modification may have species-specific function(s). As expected from figures 1-2, lactylation of some residues was induced after exposure to 25mM lactate, but inducibility varied between residues and also between species **(fig 5B).** It also varied in trophozoites versus mixed-stage cultures, implying that some residues may be lactylated stage-specifically.

**5.**
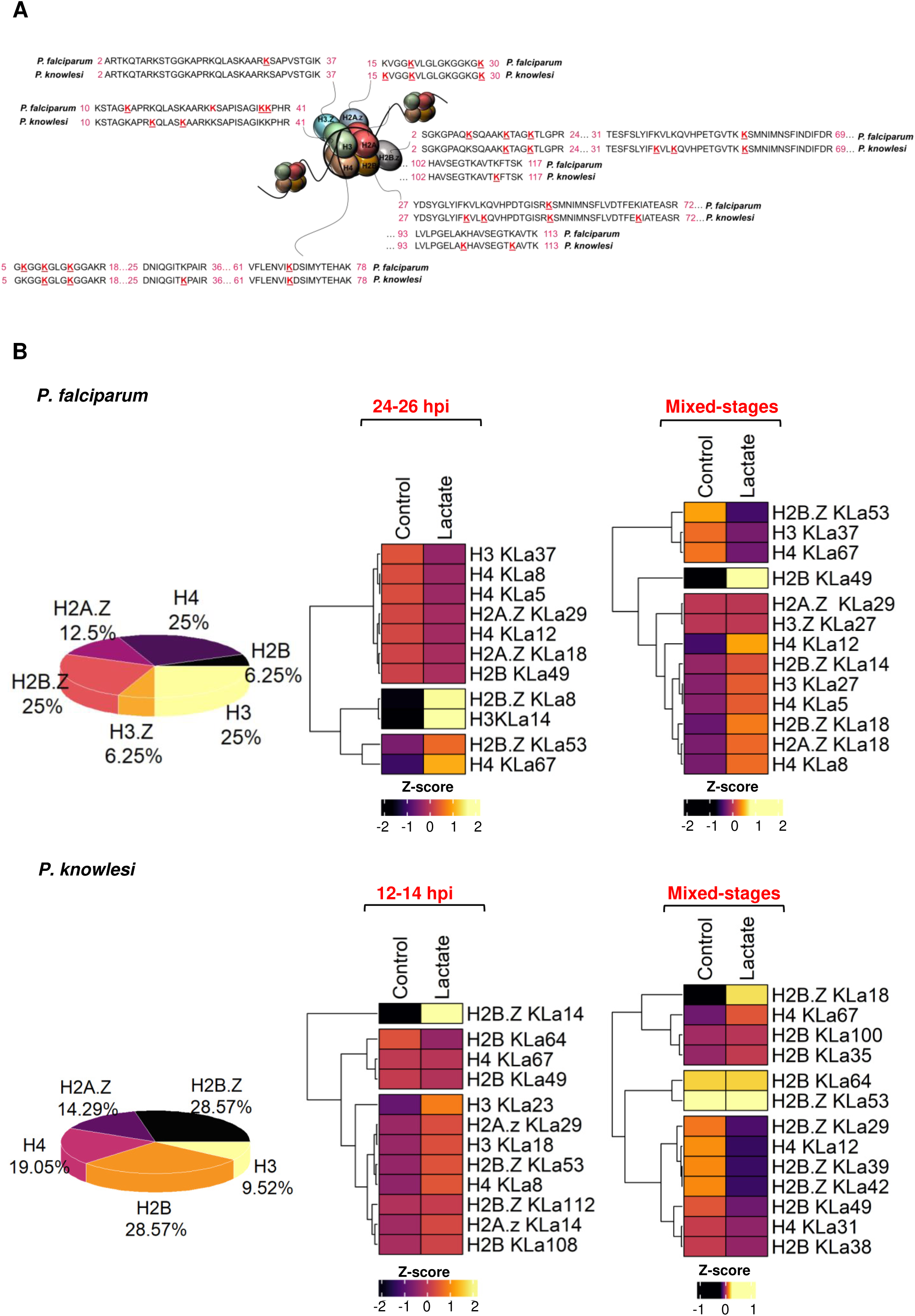
Histone lactylation occurs on multiple histone lysine residues in two *Plasmodium* species. A: Sequence alignments of histone tails in *P. falciparum* and *P. knowlesi*, showing in red all KLa sites detected by mass spectrometry (either in trophozoite or mixed-stage cultures, +/- 25mM lactate). Numbers in pink flanking the peptide sequences are residue numbers of the N- and C-termini of the peptides. The tail of histone H2A is not shown because no KLa was detected on this histone. B: Pie charts representing the distribution of KLa sites from (A), and heatmaps representing normalized intensities of KLa-containing peptides (Z-score transformed). Hierarchical clustering was used to partition the KLa sites into 3 clusters with euclidean distance and ward.D clustering algorithm. The time windows for trophozoite cultures of *P. falciparum* and *P. knowlesi* were 24-26 and 12-14 hpi, respectively.

### Cross-talk may occur between histone lactylation, acetylation and methylation

Acetylation and lactylation were often detected on the same lysine residues, both in our data and previously in human cells ^10^. Therefore we sought to investigate whether lactate exposure in the trophozoite stage could induce changes in the distribution of histone acetylation (Kac), as well as di- and tri- methylation (K(me2), K(me3)).

There was no dramatic change in the distribution of Kac sites after exposure to 25mM lactate, and Kac remained the most abundant histone modification, as previously reported ^18^ **(Tables S1-4)**. Almost all KLa sites were also detected as Kac but the converse was not true: ∼65% of Kac sites were unique **(Fig S5B)**. After lactate exposure, 44% of histone KLa sites in *P. falciparum* and 17% of those in *P. knowlesi* were significantly induced (*P*-value ≤ 0.05) **(fig 6A, top panel; Tables S1-2)**, corroborating the observation from figures 1-2 that lactylation was more strongly inducible in *P. falciparum* than in *P. knowlesi.* By contrast, only ∼12% and ∼7% of histone Kac sites, respectively, were significantly up- or down-regulated. Hence, relatively few Kac sites (amongst many that can be acetylated) must be responsible for the rise in total histone acetylation after lactate exposure, which was detected by western blot in figure 3. Overall, KLa was clearly the most inducible post-translational mofification (PTM) after lactate exposure (**fig S6A)**, and sites that were strongly induced as lactylated were not simultaneously induced as acetylated **(fig S6B)**.

**6.**
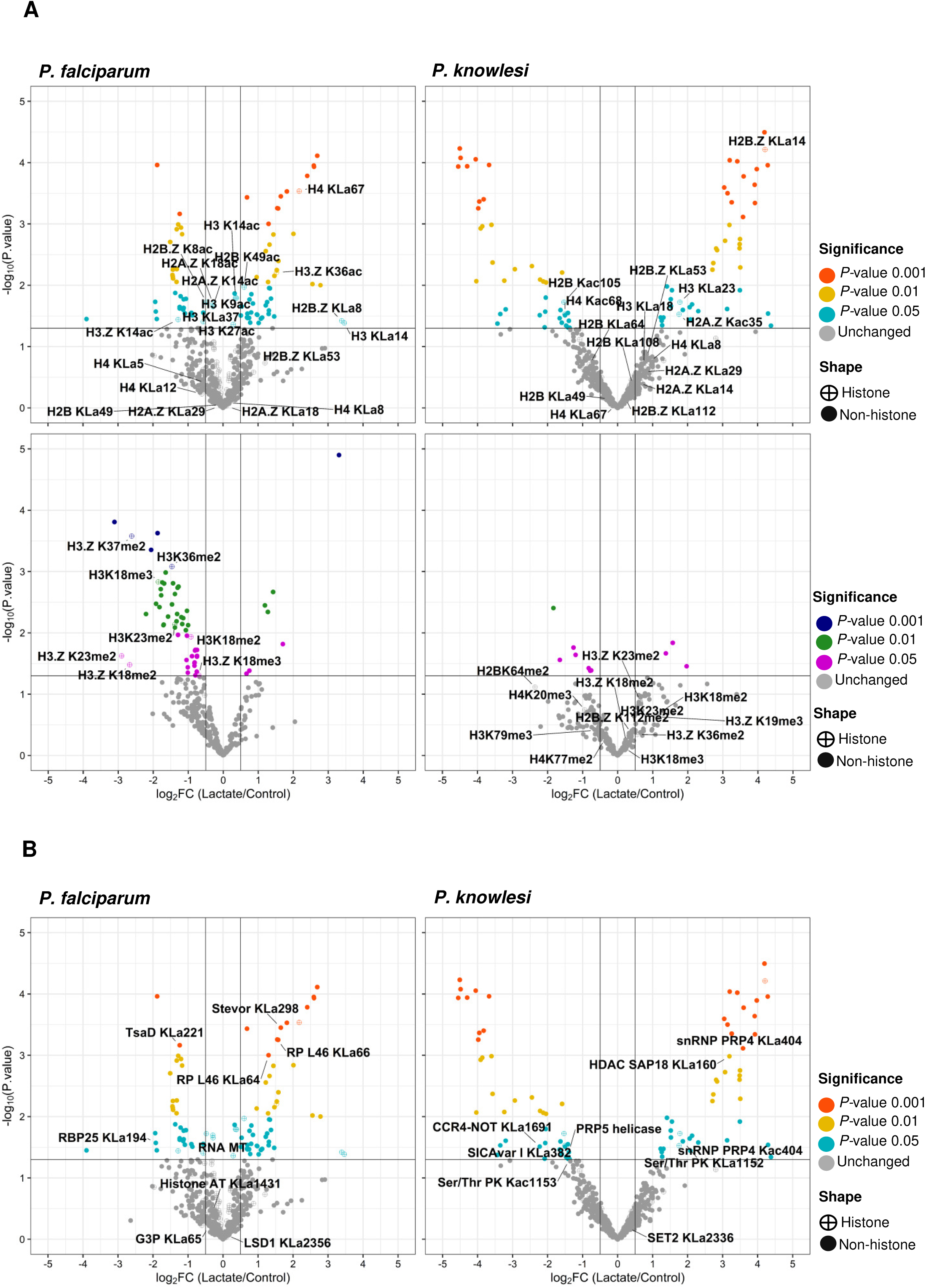
Changes in PTMs on both histones and non-histone proteins are induced by exposure to high lactate. Volcano plots representing (A) histones with differential lysine lactylation (KLa) and acetylation (Kac), in the top panels, and differential tri- or di-methylation (Kme3, Kme2) in the lower panels; (B) non-histone proteins with differential lysine lactylation (KLa). In all plots, data are from trophozoite cultures of *P. falciparum* or *P. knowlesi*, +/- 25mM lactate treatment. X-axes represent log_2_ fold changes (FC) calculated from normalized intensities, and y-axes represent *P*-values. Colored dots represent PTMs that changed significantly after lactate exposure, based on their significance threshold indicated by *P*- value. (Grey dots represent those that did not change significantly.) Symbols for histones and non-histones are hatched and filled circles, respectively. Horizontal and vertical lines represent *P*-value and log_2_FC significance thresholds. In (B), gene name abbreviations in *P. falciparum*, on the left: RNA MT, RNA methyltransferase; Histone AT, Histone acetyltransferase; LSD1, lysine specific histone demethylase; TsaD, TSaD-domain-containing protein; RBP25, RNA binding protein 25; RP L46, Ribosomal protein L46; G3P, glyceradehyde-3-phosphate dehydrogenase. In *P. knowlesi* on the right: snRNP PRP4, small nuclear ribonucleoprotein; HDAC SAP18, histone deacetylase subunit SAP18; Ser/Thr PK, serine/threonine protein kinase, CCR4-NOT, CCR4-NOT complex; PRP5 helicase, Pre-mRNA-processing ATP-dependent RNA helicase; SET2, SET2 histone methyltransferase.

In contrast to Kac, major changes did occur in K(me2) and K(me3) after lactate exposure: ∼90% of these sites were significantly down-regulated in *P. falciparum* **(fig 6A, lower panel; Tables S1-2)**. This was not the case in *P. knowlesi,* where no reciprocal changes in K(me2) and K(me3) were seen.

### Non-histone chromatin-associated proteins can be lactylated

Chromatin-associated proteins besides histones appeared in the mass spectrometry data, and many of these were also lactylated, with a subset of the modifications being induced after exogenous lactate exposure **(Fig 6B, Tables S3-4)**. In *P. falciparum*, lactylated sites were found in chromatin-bound proteins such as Alba3, several RNA helicases, proteins implicated in epigenetics (e.g. histone deacetylase 2), in glycolytic metabolism (e.g. glycerol-3-phosphate dehydrogenase) and in transcriptional control (e.g. two ApiAP2 transcription factors). A third AP2 factor, AP2G, which induces gametocytogenesis, was also detected as lactylated, but its coverage fell below the threshold for inclusion (data not shown), probably due to its very low expression in these asexual-stage cultures. An acetyl-CoA transporter was also lactylated, suggesting again the possibility of cross-talk between acetyl and lactyl epigenetic pathways **(Fig S7A; Table S3)**. In *P. knowlesi*, most of the same proteins were not detected as lactylated, although other proteins, including the acetyltransferase GCN5, were **(Fig S7B; Table S4).**

Although almost all the lactylated lysines within histones could also be acetylated **(fig S5B)**, this was not the case in non-histone proteins. Unique, lactate-induced KLa sites appeared on several chromatin-associated proteins. These included epigenetic regulators (e.g. SAP18, associated with HDAC1) and RNA binding proteins such as Alba4 in the case of *P. knowlesi*, and RNA helicases (e.g. NAM7, responsible for RNA turnover), RNA binding proteins, AP2 transcription factors and gametocytogenesis factors (e.g. Male Development 1) in the case of *P. falciparum* **(Table S1-2)**.

Collectively, it appeared that the KLa modification was widespread, both on histones and on other proteins. It was modulated on some proteins by exogenous lactate, but also probably derived from lactate produced by the parasite. The wide range of lactylated proteins suggests that the modification could regulate multiple aspects of *Plasmodium* biology, from epigenetics to RNA biology to protein function.

### Lactyl post-translational modification does not correlate with levels of protein expression

There was no consistent correlation between proteins that were inducibly modified and proteins that changed in abundance after exposure to 25mM lactate (**fig S8**). In *P. knowlesi*, no histones and very few other proteins showed significant changes (with a sca.*P*-value ≤ 0.01) after lactate exposure; in *P. falciparum*, a few proteins did change significantly in abundance, including histone H2A.Z and HP1, and interestingly the changes were mostly downregulation (**fig S8A**). Overall, however, there was no correlation between proteins whose PTMs were significantly changed, and proteins that changed in abundance (**fig S8B**). Therefore, protein lactylation does not appear to be a consistently ‘stabilizing’ nor ‘destabilizing’ PTM.

### Proteins in different biological pathways are lactylated in *P. falciparum* versus *P. knowlesi*

It was apparent that the PTMs affected by lactate exposure, both on histones and on non-histone proteins, differed in *P. falciparum* versus *P. knowlesi.* Gene ontology (GO) enrichment analysis was therefore performed to gain insight into the pathways most affected in each species.

Amongst the proteins whose PTMs changed after lactate exposure, many different biological functions were found to be enriched *(P-*value ≤ 0.05), depending on both the species and the epigenetic modification (KLa, Kac or Kme). In *P. falciparum,* lactate-modulated KLa sites were significantly enriched in proteins involved with metabolism of tRNAs and indole compounds, for example; in *P. knowlesi,* the top GO terms associated with such KLa sites included ‘cell growth’, ‘cell division’ and ‘autophagy’ **(fig 7A, fig S9, Table S5)**.

**7.**
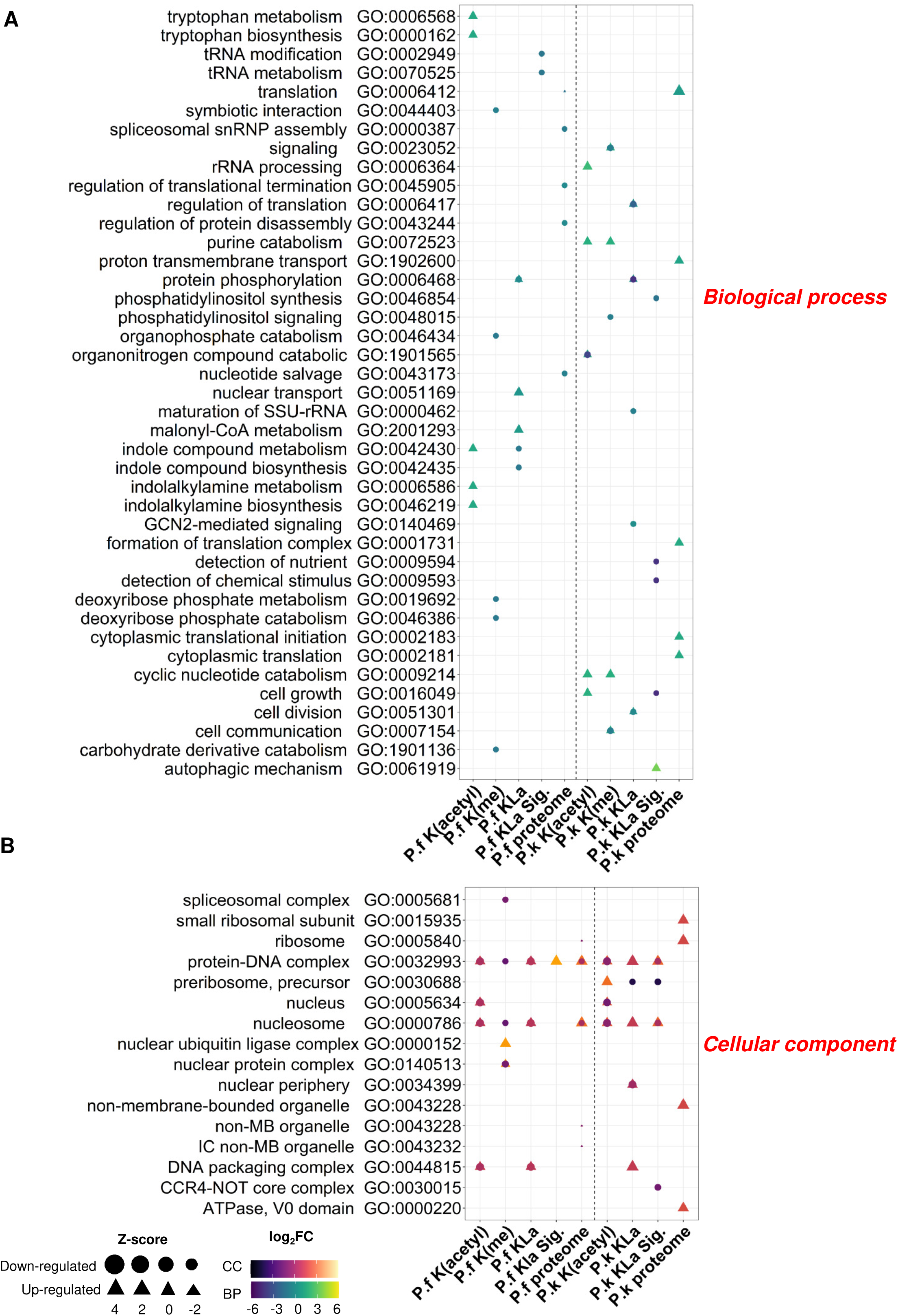
Gene ontology (GO) enrichment analysis of biological processes and cellular components enriched in PTMs after lactate exposure. Charts represent the top 5 GO terms (biological processes (A), cellular components (B)) with the highest P-value for each PTM. Triangles mark the GO terms associated with proteins whose PTMs were upregulated; circles mark those down-regulated. Shapes are color-coded based on the magnitude of the fold change. The size of the shape, represented by Z-score, indicates the magnitude of the likelihood that the term is up- or down-regulated. *P.f* and *P.k* represent *P. falciparum* (all data on the left of the vertical dashed line) and *P. knowlesi* (right of dashed line). For simplicity, K(me3) and K(me2) are both presented as K(me). ‘KLa’ refers to all KLa sites detected; KLa.Sig refers to lactylated proteins that changed significantly upon lactate exposure in trophozoites.

Amongst GO terms for cellular components, KLa sites, as well as Kac and Kme sites, that responded to lactate exposure were enriched in proteins involved with the nucleosome and protein-DNA complexes. This was not surprising given the known predominance of such PTMs on histones, as well as the location of KLa observed in figure 4, which was almost exclusively nuclear. In *P. knowlesi*, there was also an interesting enrichment of lactate-modulated KLa sites in the carbon CCR4/Not complex, which regulates aspects of protein function, RNA metabolism and transcriptional activity during late gametocyte differentiation and post-fertilization development ^19^ **(fig 7B, fig S10, Table S5).** Amongst molecular function GO terms, both species showed enrichment of ‘protein dimerization’ and ‘catalytic activity’, amongst others **(fig S11, Table S5)**.

### Blood lactate levels in malaria-infected mice correlate with distinct transcriptomic profiles in the causative parasites

In figure 1D, we investigated a rodent malaria model that becomes severely hyperlactaemic, *P. yoelii* 17XL, and showed that histone lactylation in the *P. yoelii* parasites was strongly induced in hyperlactaemic hosts. Using this model, RNAseq analysis had previously been conducted in groups of mice with early-stage versus late-stage infections (i.e. low versus high blood lactate) ^14^. We re-analysed those data for the parasite (rather than the host) transcriptome, seeking changes that correlated with high histone lactylation in late-stage infections. For comparison, we analysed the same data from *P. berghei* ANKA, where late-stage infections did not consistently generate hyperlactataemia ^14^. Rodent malarias such as these are commonly used as *in vivo* experimental systems, instead of less-accessible human malaria patients, and these existing data offered a valuable resource to suggest biological processes that might be epigenetically affected by lysine lactylation.

All datasets were first corrected for different mixes of parasite developmental stages ^20^, and indeed marked differences in the proportions of schizonts were detected in early versus late-stage *P. yoelii* infections, while gametocytes were found in low proportions in early infections but were undetectable in late infections **(Table S6).**

Differential expression (DE) analysis ^21^ performed on the corrected data revealed distinct expression profiles in late versus early stages of infection in both species **(fig S12A)**. Large cohorts of genes were differentially expressed (*P*-value ≤0.05): 502 genes in *P. yoelii* (238 upregulated and 264 downregulated) and 1521 genes in *P. berghei* (803 upregulated and 718 downregulated) (**fig S12B, Table S6**). (Using more stringent criteria, *P*-value ≤0.05 and |log2FC| ≥ 1, expression changes were similarly distributed across fewer total genes **(fig S12B)**). Late-stage *P. yoelii* parasites showed a strong downregulation of the lactate dehydrogenase (LDH) gene, as well as the glycolytic enzyme phosphofructokinase (PFK) and several genes putatively involved in gametocytogenesis. Several tRNA synthetases were also downregulated, with the interesting exception of isoleucine tRNA synthetase (isoleucine is the only amino acid that the parasite must scavenge from its host). Indeed, genes with the strongest expression changes were generally downregulated, while fewer – including several histones – were upregulated **(fig 8A,B)**. The opposite was true in late-stage *P. berghei*, where gene upregulation was most apparent, including the upregulation of phosphofructokinase and glyceraldehyde 3-phosphate dehydrogenase, and factors involved in DNA replication and repair such as MCMs and RecQ helicase **(fig 8A,B)**.

**8.**
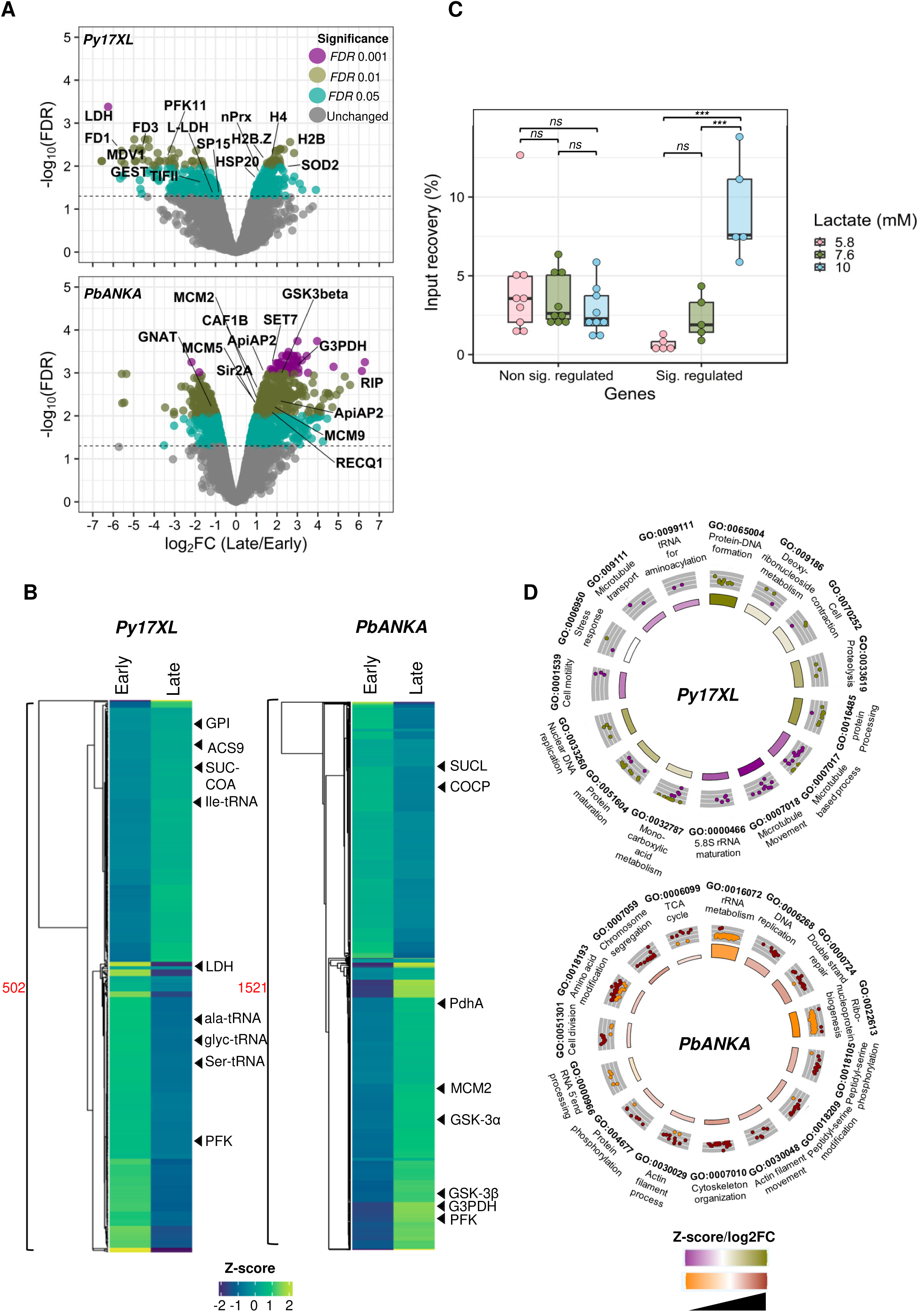
In rodent malaria parasites *in vivo,* transcriptomic profiles and lactylation of representative genes both vary with levels of host lactataemia. A: Volcano plots representing differentially expressed genes detected in late versus early infections of *P. yoelii* 17XL (top) and *P. berghei* ANKA (bottom). X-axes represent log_2_ fold changes (FC) calculated from normalized CPM values; y-axes represent *P*-values. Coloured dots represent DE genes, based on their significance threshold indicated by adjusted *P*-value (FDR) (purple dots represent those below any significance threshold). Horizontal and vertical dashed lines represent *P*-value and log_2_FC significance thresholds. Gene name abbrevations in *P. yoelli*: SP15, transcription elongation factor SPT5; TIFII, eukaryotic translation initiation factor 3 subunit L; HSP20, HSP20-like chaperone; nPrx, peroxiredoxin; PFK11, ATP-dependent 6-phosphofructokinase; H2B, histone H2B; H4, histone H4; SOD2, superoxide dismutase [Fe]; GEST, gamete egress and sporozoite traversal protein; L-LDH, L-lactate dehydrogenase; LDH, lactate dehydrogenase; FD3, female development protein FD3; H2B.Z, Histone H2B variant; MDV1, male development protein 1; FD1, female development protein 1. Gene name abbrevations in *P. berghei*: ApiAP2, AP2 domain transcription factor; GSK3beta, glycogen synthase kinase-3 beta; GNAT, N-acetyltransferase in GNAT family; MCM2/5/9, DNA replication licensing factor MCM2/5/9; RECQ1, ATP-dependent DNA helicase RecQ1; SET7, histone-lysine N-methyltransferase SET7; G3PDH, glycerol-3-phosphate dehydrogenase; RIP, PIR protein; Sir2A, histone deacetylase Sir2A; CAF1B, chromatin assembly factor 1 subunit B. B: Heatmaps representing late and early transcriptional profiles in log_2_CPM (Z-score transformed) for DE genes (502 in *P. yoelii*, 1521 in *P. berghei*). Hierarchical clustering was performed to cluster genes expression profiles with euclidean distance and ward.D clustering algorithm. C: Boxplot showing KLa enrichment detected via ChIP in the CDS of five *P. yoelii* genes whose expression significantly changed in response to hyperlactataemia (sig. regulated), and nine genes whose expression did not (non sig. regulated). Y-axis represents the percentage of chromatin input recovery (% IPed) for each ChIPed region. One biological replicate for each of three mice (i.e. three lactate conditions). Dots represent individual genes, horizontal line represents median, box represents interquartile range, whiskers represent variability outside the interquartile range (quartile -/+ 1.5 * IQR). Multiple comparison of means was performed using ANOVA and Tukey HSD post hoc tests at *p-value* < 0.05 for between-group comparison. Left boxplot (ANOVA, F = 0.788 ; d.fsum = 2; p-value = 0.466), right boxplot (ANOVA, F = 23.73; d.fsum = 2; p-value = 6.76e-05). *** p-value ≤ 0.001; ** p-value ≤ 0.01; *p-value ≤ 0.05; ns, not significant. D: Biological process gene ontology (GO) enrichment analysis of DE genes in *P. berghei* and *P. yoelii.* Plots show the top 15 GO terms with the highest *P*-values. The outer circle displays scatterplots of the expression levels (log_2_FC) for the genes (dots) in each GO term, whereas the inner circle is a bar plot where the height of the bar indicates the significance of the term (−log10 *P*-value). The colour corresponds to the Z-score, which indicates whether the term is likely to be decreased (negative value) or increased (positive value).

We used chromatin immunoprecipitation (ChIP) to check whether a selection of *P. yoelii* genes that were differentially expressed in hyperlactataemic hosts were also differentially marked with lactyl lysine **(fig 8C, fig S12C)**. Lactylation correlated strongly with blood lactate in the coding sequences of all five chosen genes, which encoded factors involved in gametocytogenesis (AP2G), metabolism (LDH, PFK), etc., whereas there was no correlation in nine control genes without differential expression. Lactylation within 1kb upstream of the same five genes was more variable and less strongly correlated with blood lactate than lactylation within the gene body (**fig S12D**). Thus, it is likely that lactylation of histones and/or other chromatin proteins within gene bodies directly regulates gene transcription, at least in the *P. yoelii* model.

When GO enrichment analysis was conducted on the up- and down-regulated genes, it was clear that late-stage *P. yoelii* parasites activated biological **(fig 8D)** and molecular **(fig S13)** functions including proteolysis and mono-carboxylic acid metabolism, while late-stage *P. berghei* parasites activated contrasting functions including DNA replication and repair, chromosome segregation, cell division and actin/cytoskeleton dynamics. Amongst functional pathways, late-stage *P. yoelii* parasites upregulated fatty acid metabolism whereas *P. berghei* upregulated sugar metabolism and glycolysis **(fig S13).** Thus, it appeared that *P. yoelii* parasites in hyperlactaemic hosts showed features of ‘dormancy’, which can be induced in *P. falciparum* by isoleucine starvation and is characterised by slowing of the replicative cycle and proteolysis ^22^; whereas *P. berghei* – in hosts without hyperlactataemia – conversely upregulated pathways involved in replication.

## DISCUSSION

This work represents the first characterisation of a new epigenetic mark, histone lactylation, in the malaria parasite *Plasmodium.* A lactylation epigenetic pathway could be highly influential in the biology and pathology of malaria, because some *Plasmodium* species, including *P. falciparum*, characteristically cause hyperlactataemia in their hosts ^3^. It would therefore make sense if these parasites had evolved to sense and respond to hyperlactataemia, potentially modulating their virulence or transmission dynamics.

Furthermore, ‘epigenetic drugs’ such as histone methyltransferase or deacetylase inhibitors have been shown to kill parasites, so epigenetic pathways in general are potential targets for antimalarial drug development ^23,24^.

Here, we show that histone lysine lactylation in several *Plasmodium* species is rapidly inducible by exogenous lactate, and also rapidly reversible. (These dynamics differ from those reported in mammalian macrophages, where histone lactylation was originally discovered, and where the modification occurred relatively slowly ^10^.) We detected histone lactylation across four *Plasmodium* species, both *in vitro* and *in vivo*, but it was not equally prominent, nor equally responsive to exogenous lactate, in all species. Rather, it was more abundant and more inducible in species that cause hyperlactataemia: *P. falciparum* and *P. yoelii*. In cultured *P. knowlesi*, a species that is only rarely associated with lactic acidosis in humans ^25^, the modification was less abundant and was not clearly regulated across a normal cell cycle, as it was in *P. falciparum.* In *P. berghei* ANKA, which did not cause consistently high levels of hyperlactataemia in mice, the modification was not clearly responsive to blood lactate, whereas in *P. yoelii* 17XL, it was. Commensurate with profound species-specific differences, mass spectrometry showed that the histone lysine residues lactylated in *P. falciparum* and *P. knowlesi* were generally different.

In human cells, it has been suggested that lactylation may act like acetylation on histone tails – as a transcription-activating mark ^10^. In *Plasmodium*, this may also be true. KLa was detected throughout nuclear chromatin, which is mainly euchromatic, with no specificity for heterochromatin. Furthermore, Kme marks are generally gene-silencing in *P. falciparum*, and mass spectrometry showed that these were mostly downregulated after lactate exposure, suggesting reciprocal regulation of KLa-induced activation and Kme-induced silencing. Again, species-specificity was apparent: such reciprocity was notably absent in *P. knowlesi*.

KLa and Kac were not, however, interchangeable: only a small subset of Kac sites in either species was detected as lactylated, and a large proportion of the KLa sites, but only a few Kac sites, were induced by exogenous lactate. This has implications for the theory that histone acetyltransferases (HATs), which normally use acetyl-CoA, can also use lactyl-CoA to act as writers of KLa ^10^. Since lysine residues were not simply modified with either moiety according to substrate abundance, our results would suggest that only some HATs might use lactyl-CoA, on only some lysine sites. In fact, if conditions of high lactate do stimulate certain HATs towards lactyltransferase activity, the same conditions appear to stimulate acetyltransferase activity on a subset of residues as well, since histone acetylation rose modestly after lactate exposure. The possible contributions of *P. falciparum* HATs to lactyltransferase activity are currently being investigated. An entirely separate pathway was also recently reported, in which alanine tRNA synthetase moonlights as a lactyltransferase ^26,27^ and this might also operate in *Plasmodium*.

Besides histone tail lactylation, we detected numerous non-histone proteins being lactylated. These were certainly not comprehensively surveyed because our mass spectrometry focussed on small, chromatin-associated proteins (large, nucleoplasmic or cytoplasmic proteins were probably under-detected). Nevertheless, we detected lactylation on ApiAP2 transcription factors, RNA processing factors, HATs and HDACs amongst others. Unlike the histone tails, where almost every lactylated lysine could also be acetylated, some non-histone proteins were uniquely lactylated (and again, they were largely species-specific between *P. falciparum* and *P. knowlesi*). Thus, the lactyl PTM may have evolved, species-specifically, to regulate *Plasmodium* biology on at least three levels: the epigenetic control of gene transcription via histone tail marks, the co-transcriptional control of RNA processing, and the post-translational modulation of protein activity in HATs, HDACs, etc. – potentially biasing their enzyme activity from acetyl- to lactyl-specificity. The biological pathways that are actually regulated await investigation, because lactylated proteins associated with a wide range of GO terms. Important pathways could potentially include stress-responsive transcriptional programmes controlled by ApiAP2s (possibly including gametocyte conversion, since lactylated ApiAP2G was detected in *P. falciparum*, albeit at a low level), and the expression of *var* genes or other virulence gene families that are epigenetically regulated via HAT/HDAC activity ^8^.

To gain insight into biological changes that can occur in parasites exposed to hyperlactatemia *in vivo*, we analysed transcriptomic data from *P. yoelii* in hyperlactaemic mice. Profound changes were detected, with features of metabolic dormancy – a programme previously reported in *P. falciparum* parasites starved of isoleucine, the only amino acid that malaria parasites cannot gain from haemoglobin catabolism ^22^. Starved *P. falciparum* parasites slowed their developmental cycle and activated proteolysis: features also seen here in the GO analysis for late-stage *P. yoelii*, but not in late-stage *P. berghei* where the hosts are not generally hyperlactaemic and the parasites do not induce high levels of histone lactylation. Five genes that were differentially expressed in *P. yoelii* during hyperlactataemia were differentially marked with lactyl lysine, strongly suggesting a direct transcription-modulating effect of lactylation – on histones and/or other chromatin proteins. Future work will focus on comprehensive analyses of the transcriptome and epigenome, to establish which transcriptional programmes in parasites exposed to hyperlactataemia are actually epigenetically controlled by histone lactylation, or by lactylation of other chromatin proteins.

## MATERIAL & METHODS

### Culture and synchronisation of *P. falciparum* and *P. knowlesi*

*Plasmodium falciparum* (NF54 strain) and *Plasmodium knowlesi* (A1-H.1 strain) were grown in RPMI 1640 media (Sigma, R4130) with 2.3 g/L sodium bicarbonate, 50 mg/L hypoxanthine (Sigma, H9377) and 25 µg/L gentamicin (Melford Laboratories, G38000-1), supplemented with 5 g/L Albumax II (Invitrogen, 11021037), 10% horse serum (Gibco, 10368902) and 4 g/L glucose for *P. knowlesi*, or with 2.5 g/L Albumax II and 5% human serum for *P. falciparum*. *P. falciparum* and *P. knowlesi* were cultured in human red blood cells (NHS Blood and Transplant) at 4% or 2% haematocrit respectively in 3% oxygen, 5% CO_2_ and 92% nitrogen gas mixture at 37°C.

For *P. falciparum* synchronization, parasites at ∼ 6% parasitaemia were initially synchronised with 5% sorbitol (w/v in water) and allowed to grow to late schizont stage. ML-10 (LifeArc) ^28,29^ was then added for 2h at a final concentration of 75 nM. Schizonts were purified via a 65% Percoll (GE Healthcare) gradient (v/v in PBS), washed twice with complete media, and allowed to reinvade in 25% haematocrit in 5 ml of complete media in the abovementioned gas mixture at 37°C for 3h with agitation at 220 rpm. 5% sorbitol was then used to remove residual schizonts and produce a tightly synchronised culture (0 to 3 hours post invasion (hpi)).

For *P. knowlesi* synchronization, parasites at ∼ 6% parasitaemia were centrifuged through a 55% Nycodenz gradient (v/v in RPMI) from 100% Nycodenz stock pH 7.5 (27.6% w/v Nycodenz, 5 mM Tris HCl, 3 mM KCl, 0.3 mM CaNa_2_·EDTA). Ring stages isolated in the bottom layer were allowed to grow to late schizont stage. ML-10 was then added to a final concentration of 150 nM for 2h. Parasites were centrifuged through a second Nycodenz gradient to collect schizonts from the top layer, which were washed twice with complete media and allowed to reinvade as above. A second Nycodenz cleanup was then performed to remove residual schizonts and produce a tightly synchronised culture (0 to 3 hpi).

### Growth of P. yoelii and P. berghei

Eight-week-old wild-type female C57BL/6J or CD1 mice were obtained from Charles River Laboratories. All mice were specified pathogen-free, housed in groups of five in individually ventilated cages, and provided ad libitum access to food and water. All experimental protocols and procedures were performed in accordance of the UK Animals (Scientific Procedures) Act (PP8697814), and were approved by the Imperial College Animal Welfare and Ethical Review Board (for *P. yoelii)* or the University of Cambridge AWERB (for *P. berghei*). The Office of Laboratory Animal Welfare Assurance for the University of Cambridge covers all Public-Health-Service-supported activities involving live vertebrates (no. A5634-01). This study was carried out in compliance with the ARRIVE guidelines (https://arriveguidelines.org/). All studies were in accordance with the Laboratory Animal Science Association’s guidelines for good practice. Prior to any experimental interventions, all mice were acclimatized to animal facilities for one week.

*P. yoelii* (17XL strain) was serially passaged through C57BL/6J mice, following previously described protocols ^14^. Blood was collected via aseptic cardiac puncture from infected donor mice under non-recovery isoflurane anesthesia and diluted in sterile phosphate-buffered saline to achieve the desired parasite concentration. Experimental mice were subsequently infected with 10^6^ live parasites via intraperitoneal (i.p.) injection.

*P. berghei* (ANKA 2.34 strain) was maintained by serial passage through CD1 mice as in previously described protocols ^30^. Experimental mice were pre-treated with 150µl of 6mg/ml phenylhydrazine i.p. three days prior to infection, to induce reticulocytosis. Blood from donor mice at 1-5% parasitaemia was collected by cardiac puncture under non-recovery Ketaset/Rompun anaesthesia and used to infected experimental mice by passage of 150µl blood, i.p..

Tail capillary blood samples were collected to prepare blood smears for parasitemia assessment and lactate measurement. Parasitemia was quantified through microscopy of thin blood smears stained with 10% Giemsa, following established protocols ^14^. Heparinized blood was collected via cardiac puncture under non-recovery anesthesia, into syringes pre-loaded with heparin, and red blood cells were separated from plasma by centrifugation at 4,000g for 10 minutes. Levels of lactate in the cardiac blood of each mouse were immediately measured using the Lactate Pro 2 meter (HAB Direct) for *P. yoelii* experiments, or the L-lactic acid (L-Lactate) assay kit (Megazyme K-LATE) for *P. berghei* experiments. Red blood cells were harvested for parasite histone extraction from ten mice per rodent malaria species, of which seven yielded a viable amount of parasite material when infected with *P. yoelii* 17XL and five yielded an appropriate amount when infected with *P. berghei* ANKA 2.34.

### Sodium L-lactate treatment of *P. falciparum* and *P. knowlesi* cultures

For titration experiments, *P. knowlesi* at 12-14 hpi and *P. falciparum* at 24-26 hpi, both at 1% parasitaemia, were washed into fresh media and then treated with 0mM, 5mM, 10mM or 25mM sodium L-lactate (Sigma), for 8h (*P. knowlesi*) and 12h (*P. falciparum*). Subsequently, samples were harvested and crude histone extractions were performed (as described below).

For timecourse experiments, *P. knowlesi* parasites at 12-14 hpi and *P. falciparum* at 24-26 hpi, both at 1% parasitaemia, were treated with 25mM L-lactate for 8h (*P. knowlesi*) and 12h (*P. falciparum*). Samples were harvested at intervals of 2-6 hours between 14-20 hpi for *P. knowlesi* and 24-36 hpi for *P. falciparum*. Lactate was then washed off at 20 hpi (*P. knowlesi*) and 36 hpi (*P. falciparum*), and fresh media was added to parasites. Samples were subsequently harvested at further intervals until reinvasion had occurred:32 hpi (for *P. knowlesi*) and 44 hpi (for *P. falciparum*). After the wash-off, media was changed every time samples were harvested. Crude histone extractions were performed from each sample. Control experiments were also performed in parallel, where no L-lactate was added, under the same conditions described above.

For mass spectrometry on histones from mixed-stage cultures of both species, parasites at a parasitaemia of ∼ 6% were washed into fresh media to remove any endogenously-produced lactate and sodium L-lactate (Sigma), dissolved in incomplete media, was added to a final concentration of 25 mM. For mass spectrometry on histones from trophozoites, *P. falciparum* at 24-26 hpi and *P. knowlesi* at 12-14 hpi were treated with sodium L-lactate. Treatments were for 12 and 9h on the two species respectively.

### Extraction of *Plasmodium* histones

*P. falciparum* and *P. knowlesi* cultures were collected by centrifugation at 800 *x g* for 5 minutes, followed by resuspending the pellet in 1.5 volumes 1X PBS. To release parasites from erythrocytes, saponin was added to a final concentration of 0.1% and incubated on ice for 10 minutes, followed by centrifugation at 12,000 *x g* at 4°C for 10 minutes. Parasite pellets were washed three times by adding ice cold 1X PBS (2 volumes of the saponin lysate) followed by centrifugation at 6000 *x g* for 2 minutes.

For *P. yoelii* and *P. berghei,* infected blood obtained by mouse cardiac puncture was placed in 50 mL RPMI HEPES modification media (‘schizont media’ - no FCS) and spun at 800 *xg* for 5 minutes, followed by an RPMI wash and centrifugation at the same speed. To remove host leukocytes, blood was passed through Plasmodipur filters (EuroProxima, 8011) using a syringe, followed by washing the filter with RPMI media to elute any remaining blood. The filtration procedure was repeated a second time before collecting the blood pellet by centrifugation at 800 *xg* for 5 minutes. Filtered blood was then mixed with 0.1 % saponin on ice for 15 minutes to release parasites, which were then collected as above.

Parasites from all species were processed either for crude or acid-based histone extraction. For crude histone extraction, parasite cell pellets were lysed by adding 1X RIPA buffer (250 mM Tris-Hcl pH 8.0, 750 mM NaCl, 5% NP-40, 5% sodium deoxycholate, 0.5% SDS, 1mM phenylmethylsulfonyl fluoride, 1X cComplete EDTA-free protease inhibitor cocktail (Roche)) followed by three freeze/thaw cycles on dry ice, 2 minutes each. Laemmli buffer was then added to a final concentration of 1X, boiled at 95 °C for 10 minutes and stored at -80°C for western blot analysis.

For mass spectrometry analysis, nuclei were liberated from parasite cell pellets by gently resuspending the parasite pellet twice in two volumes of hypotonic buffer A (10 mM Tris-HCl (pH 8.0), 3 mM MgCl₂, 0.2% v/v Nonidet P-40, 0.25 M sucrose, and a cocktail of EDTA-free protease inhibitors (Roche, 0469313200) followed by centrifugation at 4000 x *g* at 4°C for 10 minutes. The resulting chromatin pellet was then resuspended in two volumes of hypotonic buffer B (10 mM Tris-HCl, pH 8.0, 0.8 M NaCl, 1 mM EDTA [including protease inhibitor cocktail]) and incubated on ice for 10 minutes followed by centrifugation at 4000 x *g* at 4°C for 10 minutes. The chromatin pellet was then resuspended with eight volumes of 0.25 M ice-cold HCl followed by vigorous vortexing and rotation for 2h at 4 °C. Acid-soluble recovered proteins in the supernatant were collected by centrifugation at 12,000 *x g* for 30 minutes at 4 °C. Histone-containing supernatant was mixed with an equal volume of 30% TCA and incubated overnight rotating at 4 °C, followed by centrifugation at 12,000 *x g* for 15 minutes at 4 °C to collect the histone-containing pellet. This was then washed with 500 µl of ice-cold acetone and incubated at -20 °C for 3h, followed by centrifugation at 12,000 *x g* for 15 minutes. Histone pellet was then air-dried and reconstituted in 50 µl ddH_2_0. Protein concentration was measured using Qubit protein quantification kit (Invitrogen, Q33211). Purity of extracted histones was then assessed by SDS-PAGE on a 15% acrylamide gel stained with Coomassie blue G-250.

### Western blotting

Protein extracts were loaded on 15% SDS polyacrylamide gels. For lactate-titration blots, approximately 10 µg and 100 µg were loaded for *P. falciparum* and *P. knowlesi* protein extracts respectively. For timecourse blots, up to 100 µg and 200 µg were loaded for *P. falciparum* and *P. knowlesi* protein extracts respectively. For validation of histone lactylation prior to mass spectrometry, ∼ 10 µg of acid-extracted histones were used. All gels were run at 120 V for 3h and transferred to 0.45 µM nitrocellulose membrane (GE healthcare, 10600020). For both chemiluminescent (**figs 1-3**) and fluorescent (**fig S4**) blots, membranes were blocked in PBS-0.1% Tween 20 containing 5% bovine serum albumin, then probed with designated primary antibodies. For chemiluminescent detection, horseradish peroxidase (HRP)-coupled secondary antibodies were used and developed with ECL Plus Western Chemiluminescent HRP Substrate (ThermoScientific, 11527271). For fluorescent detection, fluorescent secondary antibodies were used. Blots were imaged with an Azure biosystems imager 500 and densitometry was performed with ImageJ for all western blots from titration and timecourse experiments. All lactylated histone signals were normalised using a parallel blot for total histone H4 (except for the fluorescence blot in **figure S4**, where lactylated histones were normalized using the H4 signal on the same blot). To represent data from multiple independent experiments on the same graph, the normalised values for lactylated histones within each experiment were then expressed as a fold change from the control condition (no added lactate in a titration experiment, or time-0 in a timecourse experiment).

Antibodies used in western blotting and immunofluorescence microscopy

| Primary antibody | Dilutions |  | Secondary antibody | Dilutions |  |
| --- | --- | --- | --- | --- | --- |
|  | WB | IF |  | WB | IF |
| HA - Anti-HA High Affinity Rat mAb 50 ug (Roche 11867423001) |  | 1:500 | Alexa Fluor 488-conjugated anti-rat IgG (ThermoFisher A-11006) |  | 1:1000 |
| KLA - Anti-L-Lactyllysine Rabbit mAb 100 ul (PTM-1401) | 1:1000 (chemi)<br>1:10,000 (fluor) | 1:500 | Alexa Fluor 594-conjugated anti-rabbit IgG (ThermoFisher A-11012)<br>Goat-anti-rabbit IgG IR800 (AzureSpectra, AC2134) | 1:10,000 | 1:1000 |
| H4KL12 - Anti-Lactyl-Histone H4 (Lys12) Rabbit pAB 100 ul (HOLZEL PTM-1411) | 1:000 | 1:500 | Alexa Fluor 594-conjugated anti-rabbit IgG (ThermoFisher A-11012) |  | 1:1000 |
| H3ac - Ant-Histone H3ac (pan-acetyl) Rabbit pAb 100 μg (Active motif 61637) | 1:4000 | 1:500 | Alexa Fluor 594-conjugated anti-rabbit IgG (ThermoFisher A-11012) | | 1:1000 |
| H4ac - Anti-Histone H4 (acetyl K5 + K8 + K12) 50 μg (Abcam ab233193) | 1:4000 | 1:500 | Alexa Fluor 594-conjugated anti-rabbit IgG (ThermoFisher A-11012) | | 1:1000 |
| Rabbit anti-histone H4 (Abcam, ab10158) | 1:3000 | 1:500 | HRP-conjugated goat anti-rabbit IgG (Abcam, ab6721) | 1:2500 | 1:1000 |
| Mouse anti-histone H4 (Abcam, ab17036) | 1:1000 (fluor) |  | Goat-anti-mouse IgG IR700 (AzureSpectra, AC2129) | 1:10,000 |  |

### Lactate assay

Lactate assays were performed using the L-lactic acid (L-Lactate) assay kit (Megazyme K-LATE). Media samples were taken from 5 ml cultures of *P. falciparum* NF54 and *P. knowlesi* A1-H.1, synchronised with Percoll at the ring stage, starting with 0.5% parasitaemia, every 24 h for 5 days. For two biological repeats the cultures were split every 48 h and for another two biological repeats the cultures were not split. The lactate assay was performed according to the manufacturer’s instructions (Megazyme K-LATE) on triplicate media samples of 2µl, plated in the 96-well format. The assays were read on a FLUOstar Microplate Reader (BMG LABTECH). Parasitaemia was verified by microscopy every time samples were collected. Results were analysed using GraphPad Prism.

### Immunofluorescence assay

Smears were made from *P. falciparum* 3D7 HP1_HA + BSD (gift from Prof Till Voss ^17^) and NF54, using mixed stage cultures at 4% parasitaemia. Smears were air dried, fixed in 4% paraformaldehyde/PBS for 10 minutes, washed in PBS, air dried and stored at 4°C. Parasites were permeabilised in 0.2% Triton X-100 for 15 minutes, washed in PBS, and blocked in 1% BSA/PBS with 0.1% Tween-20 for 1 h. Primary antibody labelling was done for 1 h, diluted 1:500 in 1% BSA/PBS, followed by three 5-minute washes with block (1% BSA with 0.1% Tween-20). Secondary antibody labelling was done for 1 h, diluted 1:1000 in 1% BSA/PBS, followed by one 5-minute wash with block (see table 1 for antibodies). The slides were then incubated with DAPI (4’6-diamidino-2-phenylindole) (Thermo Fisher Scientific) at 2 μg/mL in PBS for 5 minutes, followed by one 5-minute wash with block. All incubation and labelling steps were done at room temperature. Slides were cured using ProLong Diamond Antifade Mountant (Invitrogen) at room temperature and stored at 4°C prior to visualisation. Slides were imaged using a confocal fluorescence microscope Zeiss LSM700. Images were pseudo-coloured and merged using Fiji-win64 (Image J).

### Mass spectrometry

#### Sample preparation for mass spectrometry

50 µg of acid-extracted histones were resolved on SDS-PAGE 15% acrylamide gel (95 V, 3 h). Gel was stained with QC colloidal Coomassie blue G-250 (Biorad, 1610803) for 3 h, then washed with several changes of nuclease-free water over 1 h. Bands between 10 and 15

kDa were then excised and washed with several changes of 50% acetonitrile for destaining, followed by 100% acetonitrile. Gel plugs were then incubated in 0.05 M NH_4_HCO_3_ and 10 mM DTT at 56 °C for 45 minutes for disulphide bond reduction, and with 55 mM iodoacetamide and 0.1 M NH_4_ for alkylation, followed by treatment with iodoacetamide for 30 min at 30 °C in the dark. Gel plugs were then washed with water and 50% acetonitrile. For dehydration, 100% acetonitrile was added, followed by rehydration in 50 mM NH_4_HCO_3_ containing 12.5 ng/µL trypsin (Promega modified trypsin). Samples were incubated overnight at 37 °C. After digestion, the supernatant was pipetted into a sample vial and loaded onto an autosampler for automated LC-MS/MS analysis.

#### Reverse phase chromatography and mass spectrometry

All LC-MS/MS experiments were performed using a Dionex Ultimate 3000 RSLC nanoUPLC system (Thermo Fisher Scientific Inc, Waltham, MA, USA) and a Q Exactive Orbitrap mass spectrometer (Thermo Fisher Scientific Inc, Waltham, MA, USA). Separation of peptides was performed by reverse-phase chromatography at a flow rate of 300 nL/min and a Thermo Scientific reverse-phase nano Easy-spray column (Thermo Scientific PepMap C18, 2mm particle size, 100A pore size, 75 mm i.d. x 50cm length). Peptides were loaded onto a pre-column (Thermo Scientific PepMap 100 C18, 5mm particle size, 100A pore size, 300 mm i.d. x 5mm length) from the Ultimate 3000 autosampler with 0.1% formic acid for 3 minutes at a flow rate of 15 mL/min. After this period, the column valve was switched to allow elution of peptides from the pre-column onto the analytical column. Solvent A was water + 0.1% formic acid and solvent B was 80% acetonitrile, 20% water + 0.1% formic acid. The linear gradient employed was 2-40% B in 90 minutes. Further wash and equilibration steps gave a total run time of 120 minutes.

The LC eluant was sprayed into the mass spectrometer by means of an Easy-Spray source (Thermo Fisher Scientific Inc.). All m/z values of eluting ions were measured in an Orbitrap mass analyzer, set at a resolution of 35000 and was scanned between m/z 380-1500. Data dependent scans (Top 20) were employed to automatically isolate and generate fragment ions by higher energy collisional dissociation (HCD, NCE:26%) in the HCD collision cell and measurement of the resulting fragment ions was performed in the orbitrap analyser, set at a resolution of 17500. Singly charged ions and ions with unassigned charge states were excluded from being selected for MS/MS and a dynamic exclusion window of 40 was employed.

#### Data processing, protein quantification and PTM localisation

For all experiments, raw mass spectrometry data were analyzed using MaxQuant ^31^ v. 1.5.0.0 by employing Andromeda for MS/MS spectra search against the *P. falciparum* and *P. knowlesi* PlasmoDB FASTA proteome reference (release 64), and a common contaminant database ^32^. In MaxQuant, enzyme specificity was set to Trypsin/P; carbamylation of cysteines was set as a fixed modification and Acetyl lysine, lactyl lysine, and di/tri methyl lysine, as variable modifications. Missed cleavage sites were set to 5 with a maximum number of 3 modifications per peptide. For protein quantification, major protein aggregation was changed to sum. Match between run was turned on. PTMs quantification files generated by MaxQuant were exported to Perseus (v3.6.2) ^33^ to collapse localisation site information and remove reverse and contaminants. For all PTMs, sites were identified in both conditions for mixed-stage experiments, or in at least two replicates per condition for trophozoites. Remaining protein quantification and PTM quantification steps were performed in R (v3.6.2) as described below.

#### Downstream data analysis for differential protein and PTM expression

For all experiments, PTM peptide quantification files obtained after Perseus filtering steps described above, and ProteinGroup.txt protein quantification files obtained by MaxQuant, were exported to R for further processing. For mixed-stage experiments, log_2_ transformed PTM intensities were normalised using quantile normalisation using the normalizeBetweenArrays function from limma package (v3.42) ^34^. For trophozoite-stage experiments, log_2_ transformed PTM intensities were normalised using quantile and VSN normalisation for *P. falciparum* and *P. knowlesi*, respectively, using the normalizeBetweenArrays function from limma package (v3.42). VSN was chosen for the latter due to technical variability between biological replicates, which affected replicate 1 only from both + and - lactate conditions. Imputation of missing values was done by low-rank approximation of the normalised PTM data matrix using msImpute ^35^. In brief, imputation was performed with V2 method if values were missing at random, or by Barycenter approach if missingness was not at random. Differentially expressed sites were calculated using limma (two-sided, BH *p* value < 5%). At the protein level, reverse and contaminants were removed from ProteinGroup.txt quantification file, followed by extracting LFQ intensities and requiring at least two values per condition. For better accuracy in global protein levels, spectraCounteBayes function from DEqMS package (v1.22.0) ^36^ was used to make limma’s empirical-Bayesian variance calculation dependent on the number of detected peptides/PTMs per protein generated by MaxQuant, rather than estimating a fixed prior distribution for all proteins. Finally, differentially expressed proteins were calculated using limma (two-sided, BH FDR < 1%).

### RNA-seq data analysis

Raw reads from the RNA-Seq experiment generated by ^14^, belonging to Py17XL and PbANKA early and late infection stages, were downloaded, trimmed and quality checked by Trimmomatic v.0.35 ^37^. Low quality reads with Phred quality score values lower than 30 and lengths less than 20 bp were removed. (https://www.bioinformatics.babraham.ac.uk/projects/fastqc/). Filtered reads were further quality controlled by FastQC v.0.11.7 (https://www.bioinformatics.babraham.ac.uk/projects/fastqc/) and used for transcript quantification using SALMON v.0.82 ^38^. Py17XL and PbANKA reference transcript datasets were indexed using the quasi-mapping mode (--type quasi) of Salmon to create k-mers of lengths 31 (-K 31). For transcript quantification, input data sequence bias correction and bootstrapped abundance estimate were performed using the “seqBias” and “nom-Bootstraps 30” parameters from Salmon, respectively. Read counts and transcript per million reads (TPMs) were generated using tximport R package version 1.10.0 and lengthScaledTPM method ^39^ with inputs of transcript quantifications from tool SALMON. Low-expressed transcripts and genes were filtered based on analysing the data mean-variance trend. The expected decreasing trend between data mean and variance was observed when expressed transcripts were determined as those that had 3 of the 6 samples with count per million reads (CPM) 1, which provided an optimal filter for low expression. A gene was considered to be expressed if any of its transcripts with the above criteria was expressed. The TMM method was used to normalise the gene and transcript read counts to –CPM ^40^.

Prior to DE analysis and to control for heterogeneity within parasite developmental stages, we performed parasite stage deconvolution as follows. A signature matrix was taken from ^20^, which contains CPM-normalised gene expression measurements calculated across different parasite stages (male and female gametocyte, trophozoite, ring, merozoite, and schizont stages). *P. berghei* and *P. yoelii* gene names were converted to one-to-one orthologous *P. falciparum* gene names using an orthologous gene file obtained from the Malaria Cell Atlas. CPM-normalised counts alongside the signature matrix were then uploaded to CIBERSORT for deconvolution analysis using 100 permutations and disabling quantile normalisation (as recommended by CIBERSORT) to obtain the parasites cell proportions. This was performed separately for reads mapping to *P. berghei* and *P. yoelii*.

For DE analysis, limma suite was used ^34,41^ by fitting a linear model accounting for the cell proportions obtained in CIBERSORT. To compare the expression changes between conditions of experimental design, the contrast groups were set as Late-Early. For DE genes/transcript, the fold change of gene/transcript abundance were calculated based on contrast groups and significance of expression changes were determined using t-test. P-values of multiple testing were adjusted with BH to correct false discovery rate (FDR) ^42^. A gene/transcript was considered significantly DE in a contrast group if it had adjusted p-value < 0.01 and |log2FC| ≥ 1.

### Gene ontology (GO) pathway enrichment analysis

For RNA-Seq datasets, functional GO and pathway enrichment analysis were performed on significantly DE genes only, using PlasmoDB database ^43^ for annotation with default parameters. The GO terms (biological process, molecular function and cellular component) and pathways were identified to provide biological insights into the significance of DE using a *P-value* ≤ 0.05. For visualization of GO and pathways terms, combined with their associated gene expression profiles/significance, GOPlot R package v.1.02 was used ^44^.

For mass-spectrometry, functional GO enrichment analysis was performed using PlasmoDB database for annotation, with default parameters, on significantly differential proteins of the acid-extracted proteome and PTMs (for the trophozoite-stage experiment only). Exceptionally, for the KLa PTM, GO enrichment analysis was performed on both significantly and non-significantly changing lactylated proteins. Bubble plots were generated using GO terms combined with their associated genes expression profiles/significance across all PTMs and the acid-extracted proteome via in-house R scripts. For both datasets,

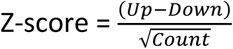

Where *up* and *down* are the number of genes up-regulated (logFC>0) in the data or down-regulated (logFC<0), respectively.

### Chromatin immunoprecipitation (ChIP) and quantitative PCR (qPCR)

Blood was harvested from three mice infected with *P. yoelii 17XL* at different levels of parasitemia and lactatataemia. Purification of erythrocytes from host leukocytes and release of parasites were performed as described above. Parasites were crosslinked immediately with 1% paraformaldehyde (Sigma) by rotating for 10 mins at 37°C, then quenching with 0.125 M glycine for 5 mins at room temperature. Nuclei were liberated by gently resuspending the parasite pellet twice in two volumes of hypotonic buffer A (10 mM Tris-HCl (pH 8.0), 3 mM MgCl₂, 0.2% v/v Nonidet P-40, 0.25 M sucrose, EDTA-free protease inhibitors (Roche, 0469313200)) and holding on ice for 15 mins. This was followed by 15 strokes using a dounce homogenizer and centrifugation at 4000 x g at 4°C for 10 mins. The chromatin pellet was then resuspended in two volumes of hypotonic buffer B (10 mM Tris-HCl, pH 8.0, 0.8 M NaCl, 1 mM EDTA, protease inhibitor cocktail) and incubated on ice for 10 mins followed by centrifugation at 4000 x g at 4°C for 10 mins. 2-3 μg of chromatin were resuspended in 50 μl of shearing buffer (1% SDS, 50 mM Tris pH 8.0, 10 mM EDTA, 1X protease inhibitor cocktail) and sheared for 47 mins using an M220 sonicator (Covaris) at 5% duty cycle, 200 cycles per burst, and 75 W of peak incident power to generate 150-400 bp fragments.

Sheared chromatin was diluted 2x in ChIP dilution buffer (0.01% SDS, 1.1% Triton X- 100, 1.2 mM EDTA pH 8.0, 25 mM Tris HCL pH 8.1, 150 mM NaCl, 1X protease inhibitor cocktail) and pre-cleared by incubating with protein A agarose beads (Thermoscientific, 20365) for 1 h at 4°C. 10% of the cleared chromatin was reserved as the input control and 90% was incubated for 4 h at 4°C using 0.75 μg of anti-KLa antibody coupled with protein A agarose beads. After one wash with low salt buffer (0.1 %SDS, 1% Triton X-100, 2 mM EDTA, 20 mM Tris-HCL pH 8.1, 150 mM NaCl), one with high salt buffer (0.1% SDS, 1% Triton X-100, 2 mM EDTA pH 8.0, 20 mM Tris-HCl pH 8.0, 500 mM NaCl), and one with lithium chloride (0.25 M LiCl, 1% NP-40, 1% sodium deoxycholate, 1 mM EDTA, 10 mM Tris-HCl pH 8.1), the immunoprecipitate was eluted twice in 85 μl elution buffer (1% SDS, 0.1M NaHCO_3_) by rotation at 65°C. ChIPed and input DNA were incubated with 2 μg RNase A (Roche, 10109169001) at 37°C for 30 min with gentle shaking. Samples were then resuspended in decrosslinking buffer (150 mM NaCl, 0.2% SDS, 600 μg Proteinase K (Ambion, AM2546)) by incubation at 65°C for 3 h with interval shaking. DNA was purified overnight at -20 °C by adding 0.1 volume of 3M sodium acetate, 4 μl of linear polyacrylamide (Ambion, AM9520) and 2.5 volumes of ice cold ethanol, and collected by centrifugation at 13,000 xg for 30 mins at 4°C. DNA pellets were washed once with 70% ice cold ethanol and resuspended in 20 μl of nuclease free water.

qPCR was conducted with 3-4 ng of DNA in technical duplicate, using a QuantStudio™ 6 Pro Real-Time PCR System: 2 mins denaturation at 95°C, 40 cycles of 5s 95°C, 30s 60°C. Data were presented as the percentage of DNA immunoprecipitated relative to input (100*2^(Ct_input_-Ct_eluates_)) . Primers were designed in the CDS and within 1kb upstream of the TSS using Primer3 and are listed below.

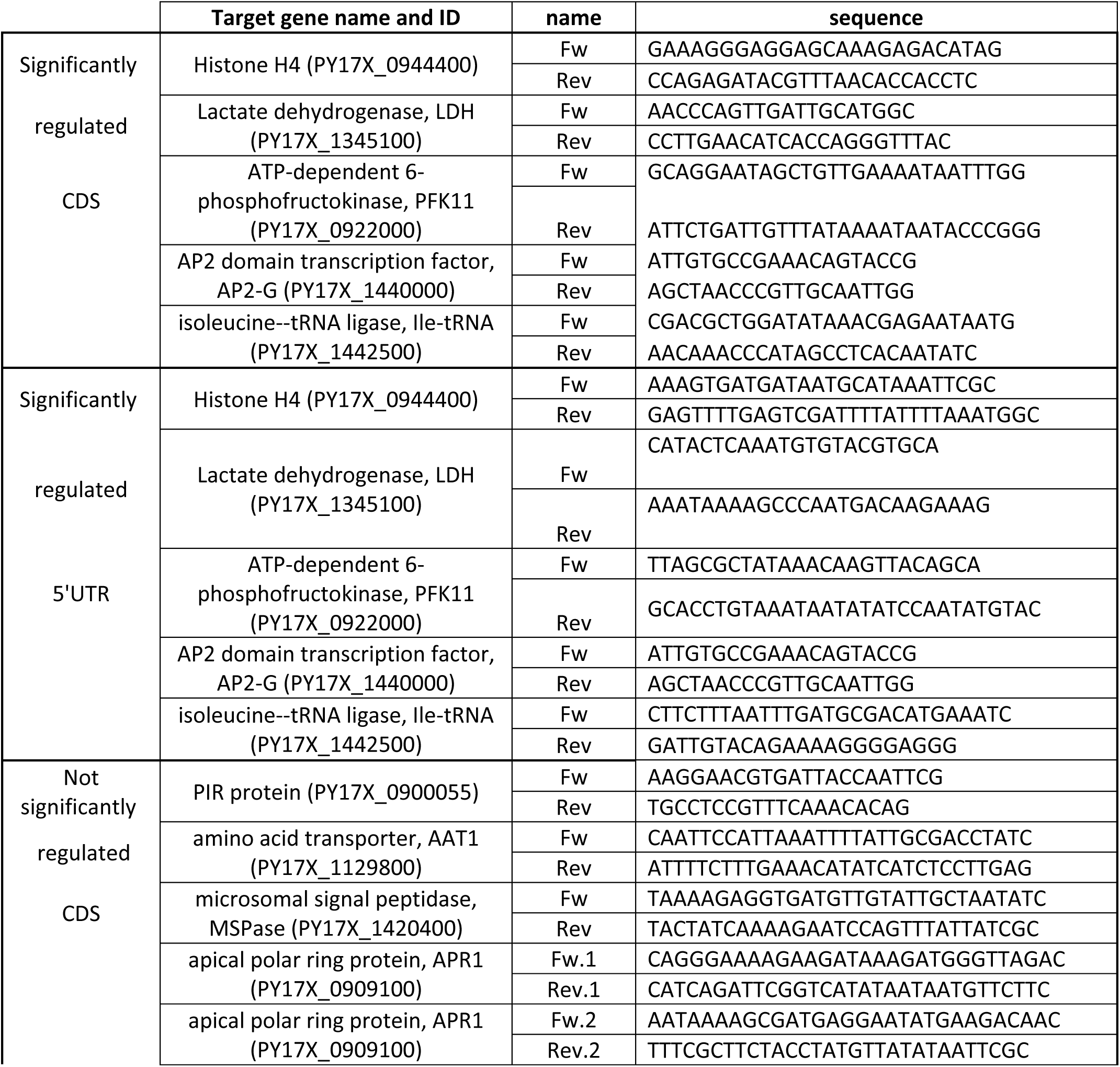
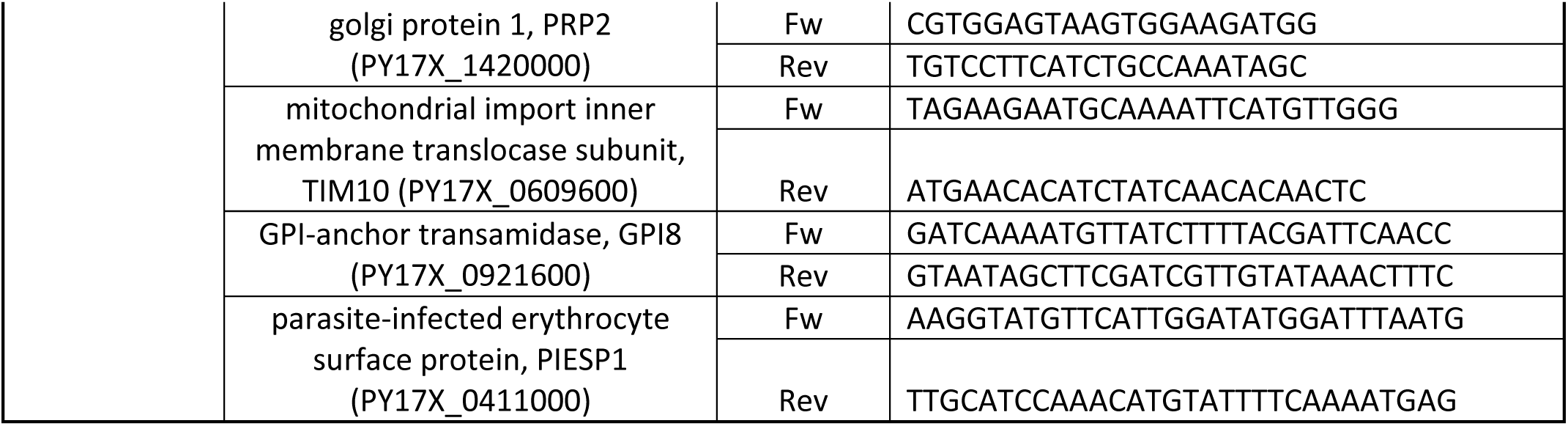

## ACKNOWLEDGEMENTS

We acknowledge the Cambridge Centre for Proteomics for mass spectrometry; the staff of Imperial College Central Biomedical Services for support with animal experiments; Till Voss (Swiss TPH) for the HP1-HA parasite line; and Rachana Ramarao and Anders Jensen for early work on this project.

## FUNDING

This work was funded by Wellcome Discovery Award 225171/Z/22/Z to CJM. AG is supported by an Imperial College Research Fellowship. AMB is funded by the MRC [MR/N00227×/1 and MR/W025701/1], Sir Isaac Newton Trust, Alborada Fund, Wellcome Trust ISSF and University of Cambridge JRG Scheme, GHIT, Rosetrees Trust (G109130) and the Royal Society (RGS/R1/201,293 and IEC/R3/19,302).

## DATA AVAILABILITY

Sequencing data from mouse, *P.yoelii* and *P. berghei* are available at ENA under the ID PRJEB43641 as per Georgiadou *et al*. ^14^. Proteome data are deposited at the ProteomeXchange Consortium via the PRIDE partner with dataset identifier PXD055236. Source data are provided with this paper.

## Notes

### Competing Interest Statement

The authors have declared no competing interest.

### Summary of Updates

Figures 1-3 and Supp data. Minor text revisions.

## REFERENCES

1 Merrick, C. J. et al. Epigenetic dysregulation of virulence gene expression in severe Plasmodium falciparum malaria. The Journal of infectious diseases 205, 1593–1600, doi:10.1093/infdis/jis239 (2012).

2 Krishna, S. et al. Lactic acidosis and hypoglycaemia in children with severe malaria: pathophysiological and prognostic significance. Transactions of the Royal Society of Tropical Medicine and Hygiene 88, 67–73, doi:10.1016/0035-9203(94)90504-5 (1994).

3 Possemiers, H., Vandermosten, L. & Van den Steen, P. E. Etiology of lactic acidosis in malaria. PLoS pathogens 17, e1009122, doi:10.1371/journal.ppat.1009122 (2021).

4 Andrade, C. M. et al. Increased circulation time of Plasmodium falciparum underlies persistent asymptomatic infection in the dry season. Nature medicine 26, 1929–1940, doi:10.1038/s41591-020-1084-0 (2020).

5 Mancio-Silva, L. et al. Nutrient sensing modulates malaria parasite virulence. Nature 547, 213–216, doi:10.1038/nature23009 (2017).

6 Harris, C. T. et al. Sexual differentiation in human malaria parasites is regulated by competition between phospholipid metabolism and histone methylation. Nat Microbiol 8, 1280–1292, doi:10.1038/s41564-023-01396-w (2023).

7 Diffendall, G. et al. RNA polymerase III is involved in regulating Plasmodium falciparum virulence. eLife 13, doi:10.7554/eLife.95879.2 (2024).

8 Duraisingh, M. T. & Horn, D. Epigenetic Regulation of Virulence Gene Expression in Parasitic Protozoa. Cell host & microbe 19, 629–640, doi:10.1016/j.chom.2016.04.020 (2016).

9 West, R. & Sullivan, D. J. Lactic acid supplementation increases quantity and quality of gametocytes in Plasmodium falciparum culture. Infection and immunity 15, e00635–00620, doi:10.1128/IAI.00635-20 (2020).

10 Zhang, D. et al. Metabolic regulation of gene expression by histone lactylation. Nature 574, 575–580, doi:10.1038/s41586-019-1678-1 (2019).

11 Dai, E., Wang, W., Li, Y., Ye, D. & Li, Y. Lactate and lactylation: Behind the development of tumors. Cancer Lett 591, 216896, doi:10.1016/j.canlet.2024.216896 (2024).

12 Su, J. et al. Functions and mechanisms of lactylation in carcinogenesis and immunosuppression. Front Immunol 14, 1253064, doi:10.3389/fimmu.2023.1253064 (2023).

13 Merrick, C. J. Histone lactylation: a new epigenetic axis for host-parasite signalling in malaria? Trends Parasitol 39, 12–16, doi:10.1016/j.pt.2022.10.004 (2023).

14 Georgiadou, A. et al. Comparative transcriptomic analysis reveals translationally relevant processes in mouse models of malaria. Elife 11, doi:10.7554/eLife.70763 (2022).

15 Hikosaka, K., Hirai, M., Komatsuya, K., Ono, Y. & Kita, K. Lactate retards the development of erythrocytic stages of the human malaria parasite Plasmodium falciparum. Parasitol Int 64, 301–303, doi:10.1016/j.parint.2014.08.003 (2015).

16 Duffy, S. & Avery, V. M. Routine In Vitro Culture of Plasmodium falciparum: Experimental Consequences? Trends Parasitol 34, 564–575, doi:10.1016/j.pt.2018.04.005 (2018).

17 Flueck, C. et al. Plasmodium falciparum heterochromatin protein 1 marks genomic loci linked to phenotypic variation of exported virulence factors. PLoS pathogens 5, e1000569 (2009).

18 Trelle, M. B., Salcedo-Amaya, A. M., Cohen, A. M., Stunnenberg, H. G. & Jensen, O. N. Global histone analysis by mass spectrometry reveals a high content of acetylated lysine residues in the malaria parasite Plasmodium falciparum. J Proteome Res 8, 3439–3450, doi:10.1021/pr9000898 (2009).

19 Hart, K. J., Power, B. J., Rios, K. T., Sebastian, A. & Lindner, S. E. The Plasmodium NOT1-G paralogue is an essential regulator of sexual stage maturation and parasite transmission. PLoS biology 19, e3001434, doi:10.1371/journal.pbio.3001434 (2021).

20 Tebben, K., Dia, A. & Serre, D. Determination of the Stage Composition of Plasmodium Infections from Bulk Gene Expression Data. mSystems 7, e0025822, doi:10.1128/msystems.00258-22 (2022).

21 Calixto, C. P. G. et al. Rapid and Dynamic Alternative Splicing Impacts the Arabidopsis Cold Response Transcriptome. Plant Cell 30, 1424–1444, doi:10.1105/tpc.18.00177 (2018).

22 Babbitt, S. E. et al. Plasmodium falciparum responds to amino acid starvation by entering into a hibernatory state. Proceedings of the National Academy of Sciences of the United States of America 109, E3278–3287, doi:10.1073/pnas.1209823109 (2012).

23 Wang, M. et al. Drug Repurposing of Quisinostat to Discover Novel Plasmodium falciparum HDAC1 Inhibitors with Enhanced Triple-Stage Antimalarial Activity and Improved Safety. Journal of medicinal chemistry 65, 4156–4181, doi:10.1021/acs.jmedchem.1c01993 (2022).

24 Malmquist, N. A., Moss, T. A., Mecheri, S., Scherf, A. & Fuchter, M. J. Small-molecule histone methyltransferase inhibitors display rapid antimalarial activity against all blood stage forms in Plasmodium falciparum. Proceedings of the National Academy of Sciences of the United States of America 109, 16708–16713, doi:10.1073/pnas.1205414109 (2012).

25 Barber, B. E., Grigg, M. J., William, T., Yeo, T. W. & Anstey, N. M. The Treatment of Plasmodium knowlesi Malaria. Trends Parasitol 33, 242–253, doi:10.1016/j.pt.2016.09.002 (2017).

26 Ju, J. et al. The alanyl-tRNA synthetase AARS1 moonlights as a lactyltransferase to promote YAP signaling in gastric cancer. J Clin Invest 134, doi:10.1172/JCI174587 (2024).

27 Zong, Z. et al. Alanyl-tRNA synthetase, AARS1, is a lactate sensor and lactyltransferase that lactylates p53 and contributes to tumorigenesis. Cell 187, 2375–2392 e2333, doi:10.1016/j.cell.2024.04.002 (2024).

28 Large, J. M. et al. Potent bicyclic inhibitors of malarial cGMP-dependent protein kinase: approaches to combining improvements in cell potency, selectivity and structural novelty. Bioorg Med Chem Lett 29, 126610, doi:10.1016/j.bmcl.2019.08.014 (2019).

29 Large, J. M. et al. Potent inhibitors of malarial P. Falciparum protein kinase G: Improving the cell activity of a series of imidazopyridines. Bioorg Med Chem Lett 29, 509–514, doi:10.1016/j.bmcl.2018.11.039 (2019).

30 Blagborough, A. M. et al. Transmission-blocking interventions eliminate malaria from laboratory populations. Nature communications 4, 1812, doi:10.1038/ncomms2840 (2013).

31 Cox, J. & Mann, M. MaxQuant enables high peptide identification rates, individualized p.p.b.- range mass accuracies and proteome-wide protein quantification. Nat Biotechnol 26, 1367–1372, doi:10.1038/nbt.1511 (2008).

32 Cox, J. et al. Andromeda: a peptide search engine integrated into the MaxQuant environment. J Proteome Res 10, 1794–1805, doi:10.1021/pr101065j (2011).

33 Tyanova, S. et al. The Perseus computational platform for comprehensive analysis of (prote)omics data. Nature methods 13, 731–740, doi:10.1038/nmeth.3901 (2016).

34 Ritchie, M. E. et al. limma powers differential expression analyses for RNA-sequencing and microarray studies. Nucleic acids research 43, e47, doi:10.1093/nar/gkv007 (2015).

35 Hediyeh-Zadeh, S., Webb, A. I. & Davis, M. J. MsImpute: Estimation of Missing Peptide Intensity Data in Label-Free Quantitative Mass Spectrometry. Mol Cell Proteomics 22, 100558, doi:10.1016/j.mcpro.2023.100558 (2023).

36 Zhu, Y. et al. DEqMS: A Method for Accurate Variance Estimation in Differential Protein Expression Analysis. Mol Cell Proteomics 19, 1047–1057, doi:10.1074/mcp.TIR119.001646 (2020).

37 Bolger, A. M., Lohse, M. & Usadel, B. Trimmomatic: a flexible trimmer for Illumina sequence data. Bioinformatics 30, 2114–2120, doi:10.1093/bioinformatics/btu170 (2014).

38 Patro, R., Duggal, G., Love, M. I., Irizarry, R. A. & Kingsford, C. Salmon provides fast and bias-aware quantification of transcript expression. Nature methods 14, 417–419, doi:10.1038/nmeth.4197 (2017).

39 Soneson, C., Matthes, K. L., Nowicka, M., Law, C. W. & Robinson, M. D. Isoform prefiltering improves performance of count-based methods for analysis of differential transcript usage. Genome biology 17, 12, doi:10.1186/s13059-015-0862-3 (2016).

40 Bullard, J. H., Purdom, E., Hansen, K. D. & Dudoit, S. Evaluation of statistical methods for normalization and differential expression in mRNA-Seq experiments. BMC bioinformatics 11, 94, doi:10.1186/1471-2105-11-94 (2010).

41 Law, C. W., Chen, Y., Shi, W. & Smyth, G. K. voom: Precision weights unlock linear model analysis tools for RNA-seq read counts. Genome biology 15, R29, doi:10.1186/gb-2014-15-2-r29 (2014).

42 Benjamini, Y. & Yekutieli, D. Quantitative trait Loci analysis using the false discovery rate. Genetics 171, 783–790, doi:10.1534/genetics.104.036699 (2005).

43 Aurrecoechea, C. et al. PlasmoDB: a functional genomic database for malaria parasites. Nucleic acids research 37, D539–543, doi:10.1093/nar/gkn814 (2009).

44 Walter, W., Sanchez-Cabo, F. & Ricote, M. GOplot: an R package for visually combining expression data with functional analysis. Bioinformatics 31, 2912–2914, doi:10.1093/bioinformatics/btv300 (2015).

