## Supplementary figures and legends for "Histone lactylation: a new epigenetic mark in the malaria parasite *Plasmodium*"

**SUPPLEMENTARY FIGURE LEGENDS**

1. *Plasmodium berghei* histones are not inducibly lactylated in hyperlactaemic mice
   Lysine lactylation was measured in *P. berghei* ANKA parasites exposed to varying levels of lactate in the blood of the host mouse. Western blots show 'Pan KLa' and histone H4 as a control. M1-M5, individual mice; L, ladder. Scatter plot represents Pearson correlation ($R^2$= 0.06) between histone KLa signal intensities normalized to H4 (*y*-axis) and lactate levels in the blood (*x*-axis) upon parasite collection from host.

2. The effect upon *P. falciparum* cell-cycle progression of treatment with 25mM lactate for 12h

A: *P. falciparum* parasitaemia, blind-counted by microscopy, at each timepoint of the two timecourse experiments shown in figure 2A. Parasites were treated with 25mM lactate for 12h from the early trophozoite stage (or were untreated in the control); then in both conditions parasites were washed and regular counts were continued to the point of complete reinvasion in the subsequent cycle. The mean and range of counts from two independent timecourses are shown; n ≥ 20 parasites per timepoint. At 24h, a slight delay in reinvasion was apparent in the two lactate-treated timecourses, but reinvasion rates recovered to control levels by 36h.

B: Staging of parasites by microscopy (n ≥ 20 per timepoint) for the timecourses in (A). ET, early trophozoite; MT, mid trophozoite; LT, late trophozoite; S, schizont; R, ring. A slight delay in progression of mature stages (i.e. late trophozoites and schizonts) in the two lactate-treated timecourses is apparent here, as it is in (A), but homogenous populations of rings appeared in both timecourses, +/- 25mM lactate, by 36h.

3. Histone lactylation is distributed throughout the nucleus

A: Confocal microscopy images showing the nuclear locations of acetyl histone H4 (H4ac) and of HP1 in *P. falciparum* 3D7 HP1_HA. H4ac, red; HP1_HA, green; DAPI, blue.

B: Confocal microscopy images showing the location of KLa in *P. falciparum* NF54. This confirms that the location is the same as that shown in main figure 4, and is not affected by the tagging of HP1 in the line 3D7 HP1_HA (strain used in main figure 4). KLa, red; DAPI, blue.

C: Images as in (B), showing lactyl-H4K12 in the *P. falciparum* NF54 strain.

D: Images as in (B), showing H4ac in the *P. falciparum* NF54 strain.

Scale bar (2 µm) applies to all images. R, ring stage, T, trophozoite stage; S, schizont stage.

4. Acid based histone extraction: quality control

A: Coomassie-stained gel of ~ 25 µg of acid-extracted histones with (**+**) /without (**-**) lactate (lac) resolved between ~ 10 and 15 kDa. Molecular weight marker (MW) in kDa is indicated to the left. Histones and histone variants names in *Plasmodium* species are indicated to the right.

B: Immunoblots showing differential lysine lactylation of histones from synchronized trophozoites in response to 25 mM lactate in *P. knowlesi* and *P. falciparum* (biological triplicates). In each gel, lanes marked (+) and (-) correspond to ~ 1 ug of acid-extracted histones with or without lactate treatment. H4 is used as a loading control. Quantified signals are displayed to the right, as bar graphs. Bars show the mean band intensity from biological replicates, normalized to H4. Error bars show the standard error of the mean. T-tests were performed to check for significant changes between groups (0 mM versus 25 mM lactate), n=3/group, *** indicates *P*-value ≤ 0.01.

5. <u>Distribution of lysine lactylation: peptide intensities</u>

A) Violin plot representing lactylated peptide intensities detected by mass spectrometry for samples with (Lactate) and without (Control) 25mM lactate treatment, with the middle horizontal line representing the median. The yellow shape shows the distribution of the data, and the grey dots represent the $\log_2$ transformed lactylation values for individual peptides.

B) Venn diagram representing histone lysine (K) sites of *Plasmodium* that can be either uniquely lactylated (KLa) (dark pink), acetylated (Kac) (yellow), or lactylated and acetylated interchangeably (light pink).

6. <u>KLa profiles on histones are more inducible after lactate exposure than other PTMs</u>

A) Heatmap representing normalized intensities of significantly differential PTM sites (Z-score transformed) after 25mM lactate treatment: histone lysine lactylation (KLa), acetylation (Kac), trimethylation (Kme3) and dimethylation (Kme2). Hierarchical clustering was performed for each species, *P. falciparum* and *P. knowlesi,* independently to cluster the PTM site expression profiles with euclidean distance and ward.D clustering algorithm. The colored bar on the left represents the categories of histone PTM (lactyl/acetyl/methyl).

B) Heatmap representing normalized intensities of the few histone Kac sites that were significantly differential, and that can be interchangeably – and significantly differentially – lactylated as well, in response to 25 mM lactate in *P. falciparum*.

7. <u>KLa profiles in the acid-extracted proteome of *P. knowlesi* and *P. falciparum*</u>

A) Heatmap representing normalized peptide intensities of KLa sites on histone (red) and non-histone (black) proteins (Z-score transformed). Data from *P. falciparum* mixed-stage culture, +/-25 mM lactate treatment. Hierarchical clustering was performed to cluster protein expression profiles with euclidean distance and ward.D clustering algorithm.

B) Heatmap representing normalized peptide intensities of KLa sites on histone (red) and non-histone (black) proteins (Z-score transformed). Data from *P. knowlesi* mixed-stage culture, +/-25 mM lactate treatment. Hierarchical clustering was performed to cluster protein expression profiles with euclidean distance and ward.D clustering algorithm.

8. <u>Comparison between the *Plasmodium* proteome and lactate-induced PTM landscape shows little correlation between PTMs and protein abundance</u>

A) Volcano plots representing differential proteome changes in *Plasmodium* trophozoites in response to 25 mM lactate treatment. X-axes represent $\log_2$ fold changes (FC) calculated from normalized intensities, and y-axes represents $-\log_{10}$ transformed scaled adjusted *P*-values, and scaled *P*-values, for *P. falciparum and P. knowlesi*, respectively. Colored dots represent significantly changing proteins, based on their significance threshold indicated by *P*-value. Dashed black horizontal and vertical lines represent *P*-value and $\log_2$FC significance thresholds.

B) Scatter plot representing correlation between significantly changing lactylated peptides and their protein abundance. $\log_2$FC of the lactylated peptide is plotted on the x-axis and the corresponding proteome $\log_2$FC is plotted on the y-axis. Pearson correlation coefficient (Pearson R) between both entities for each *Plasmodium* species is shown in red. Histone and non-histone proteins are in light orange and sky-blue, respectively. The linear regression fit is shown by the green line (*x* and *y* reflect $\log_2$FC on x-axes and y-axes, respectively). Only proteins with 2 unique razor+ peptides were included in the analysis.

9. <u>GO enrichment analysis of biological processes related to PTMs that are influenced by lactate exposure</u>

GO terms are shown for the proteins with significantly differential PTMs, and for proteins that changed significantly in the chromatin-associated proteome, after 25mM lactate treatment. Each GO term is represented with the genes associated with it as up- (triangle) or/and down- (circle) regulated. Shapes are color-coded based on the magnitude of the fold change. The size of the shape, represented by Z-score, indicates whether the term is likely to be decreased (negative value) or increased (positive value). For simplicity, K(me3) and K(me2) are presented as K(me). *P.f* and *P.k* represent Plasmodium *knowlesi* and *falciparum*, respectively. KLa.Sig and KLa represent GO terms enrichment for differentially expressed KLa proteins and lactylated acid-extracted proteome, respectively. Dashed vertical lines separate *P. knowlesi* and *P. falciparum* GO enrichment.

10. <u>GO enrichment analysis of cellular components related to PTMs that are influenced by lactate exposure</u>

GO terms are shown for the proteins with significantly differential PTMs, and for proteins that changed significantly in the chromatin-associated proteome, after 25mM lactate treatment. Each GO term is represented with the genes associated with it as up- (triangle) or/and down- (circle) regulated. Shapes are color-coded based on the magnitude of the fold change. The size of the shape, represented by Z-score, indicates whether the term is likely to be decreased (negative value) or increased (positive value). For simplicity, K(me3) and K(me2) are presented as K(me). *P.f* and *P.k* represent

Plasmodium *knowlesi* and *falciparum*, respectively. KLa.Sig and KLa represent GO terms enrichment for differentially expressed KLa proteins and lactylated acid-extracted proteome, respectively. Dashed vertical lines separate *P. knowlesi* and *P. falciparum* GO enrichment.

11. <u>GO enrichment analysis of molecular function related to PTMs that are influenced by lactate exposure</u>

GO terms are shown for the proteins with significantly differential PTMs, and for proteins that changed significantly in the chromatin-associated proteome, after 25mM lactate treatment. Each GO term is represented with the genes associated with it as up- (triangle) or/and down- (circle) regulated. Shapes are color-coded based on the magnitude of the fold change. The size of the shape, represented by Z-score, indicates whether the term is likely to be decreased (negative value) or increased (positive value). For simplicity, K(me3) and K(me2) are presented as K(me). *P.f* and *P.k* represent Plasmodium *knowlesi* and *falciparum*, respectively. KLa.Sig and KLa represent GO terms enrichment for differentially expressed KLa proteins and lactylated acid-extracted proteome, respectively. Dashed vertical lines separate *P. knowlesi* and *P. falciparum* GO enrichment.

12. <u>Differential gene expression and KLa chromatin enrichment profiles in response to hyperlactataemia</u>

A) Principal component analysis of RNA-Seq gene expression data representing normalized counts of 4983 and 4712 genes from three biological replicates (Rn) corresponding to *P. yoelii* (top panel) and *P. berghei* (bottom panel), respectively, at early (pink), and late (green) stages of mouse infection.

B) Violin plot representing the distribution and number of significantly up- and down-regulated genes (*P*-value ≤0.05 (Top panel) or *P*-value ≤0.01 and |log2FC| ≥ 1 (lower panel) in *P. yoelii* 17XL (green) and *P. berghei* ANKA (purple) in late-stage versus early-stage infection . Violin width is irrespective of data range.

C) KLa enrichment in the CDS of 5 differentially expressed genes (upper panel) and 9 control genes (lower panel) in *P. yoelii* after varying lactate exposure. Barplots show the percentage of chromatin input recovery (% IPed) for one region of each gene.

D) KLa enrichment upstream of 5 differentially expressed genes, as in (C). Upper panel: Y-axis represents the percentage of chromatin input recovery (% IPed) for each ChIPed region. One biological replicate for each of three mice (i.e. three lactate conditions). Dots represent individual genes, horizontal line represents median, box represents interquartile range, whiskers represent variability outside the interquartile range (quartile -/+ 1.5 * IQR). Multiple comparison of means was performed using ANOVA and Tukey HSD post hoc tests at p-value < 0.05 for between-group comparaison. ANOVA, F = 4.413 ; d.fsum = 2; p-value = 0.0366). *** p-value ≤ 0.001; ** p-value ≤ 0.01; *p-value ≤ 0.05; ns, not significant. Lower panel: barplot as in (C).

13. GO enrichment analysis of genes differentially expressed at late versus early stage of infection in *P.yoelii* 17XL and *P.berghei* ANKA

A) Molecular function gene ontology (GO) enrichment analysis of DE genes in *P. berghei* and *P. yoelii*. Plots show the top 15 GO terms enriched in molecular function.

B) Top 15 GO terms enriched in cellular components of DE genes in *P. berghei* and *P. yoelii*.

C) Top 15 pathways terms enriched in DE genes in *P. berghei* and *P. yoelii*.

For all plots the outer circle displays scatterplots of the expression levels (log2FC) for the genes (dots) in each GO term, whereas the inner circle is a bar plot where the height of the bar indicates the significance of the term (−log10 P-value), and color corresponds to the Z-score, which indicates if the term is likely to be decreased (negative value) or increased (positive value).

### SUPPLEMENTARY TABLE LEGENDS

Supplementary Table 1. *P. falciparum* trophozoite mass spectrometry and differential expression analysis

Supplementary Table 2. *P. knowlesi* trophozoite mass spectrometry and differential expression analysis

Supplementary Table 3. *P. falciparum* mixed stages mass spectrometry analysis

Supplementary Table 4. *P. knowlesi* mixed stages mass spectrometry analysis

Supplementary Table 5. Gene ontology of differentially expressed PTMs sites and acid-extracted proteome components in *P. falciparum* and *P. knowlesi* trophozoites.

Supplementary Table 6. RNA-Seq analysis and GO enrichment of DE genes in *P. berghei* and *P. yoelii.*

Supplementary Figure 1

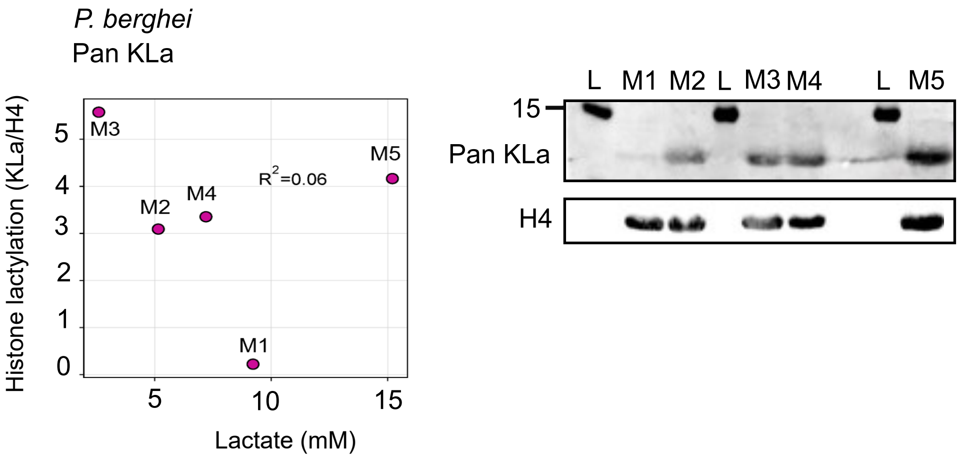

Supplementary Figure 2

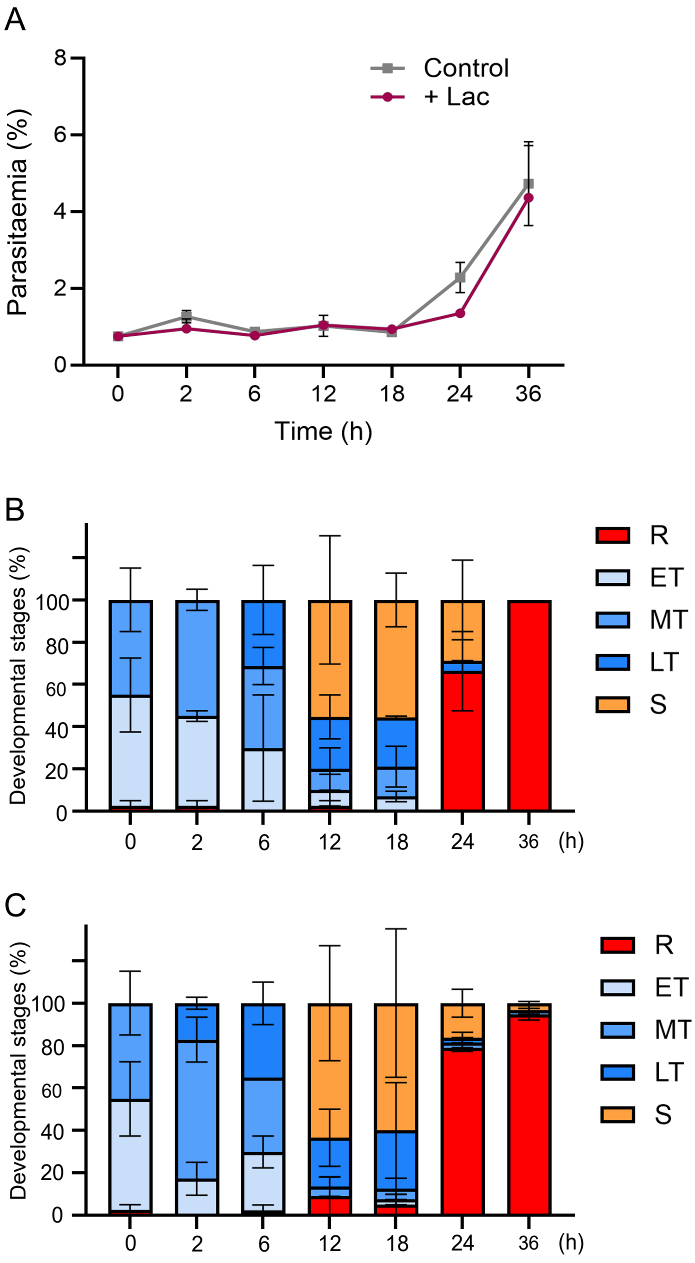

Supplementary figure 3

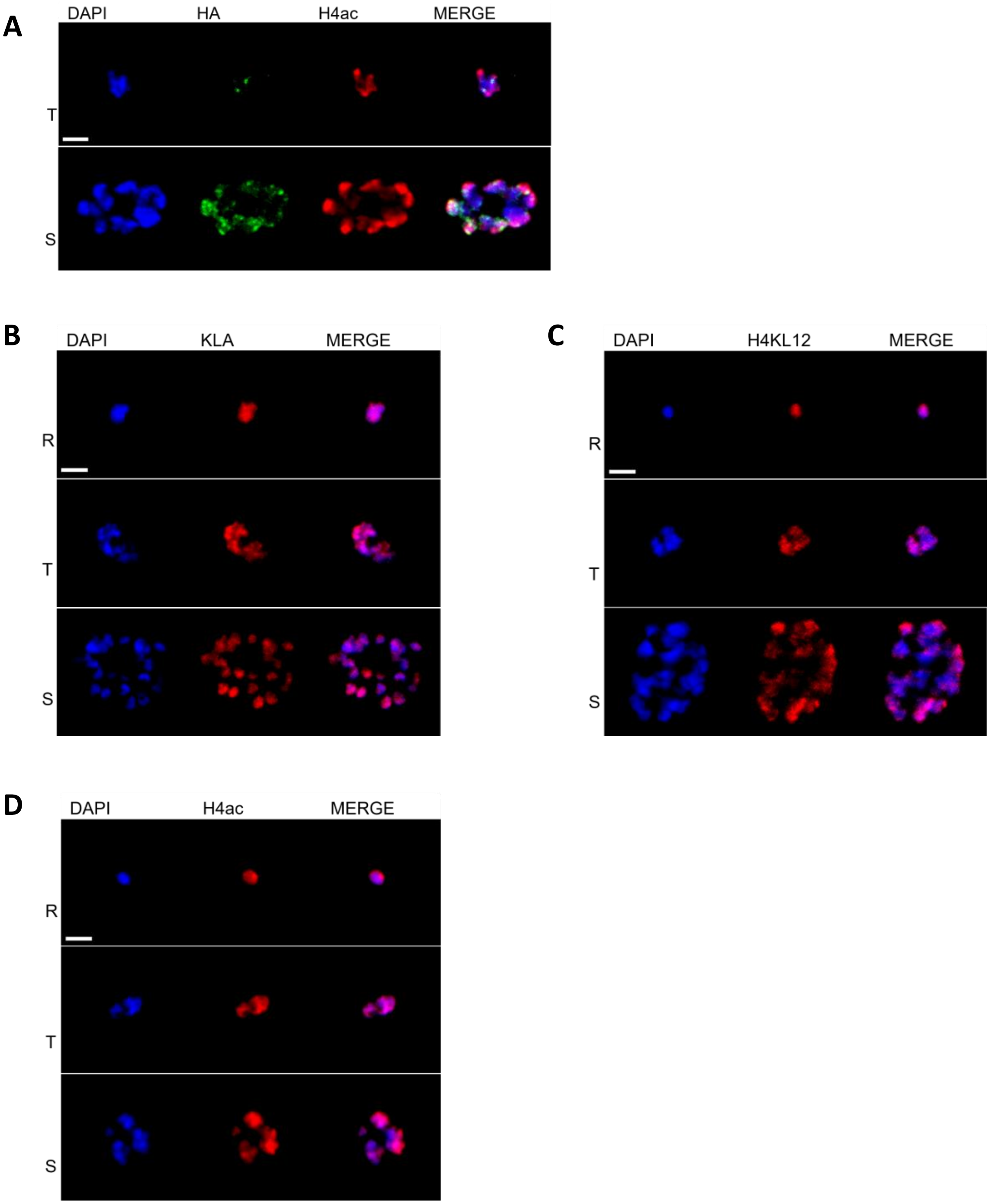

Supplementary figure 4

**A**

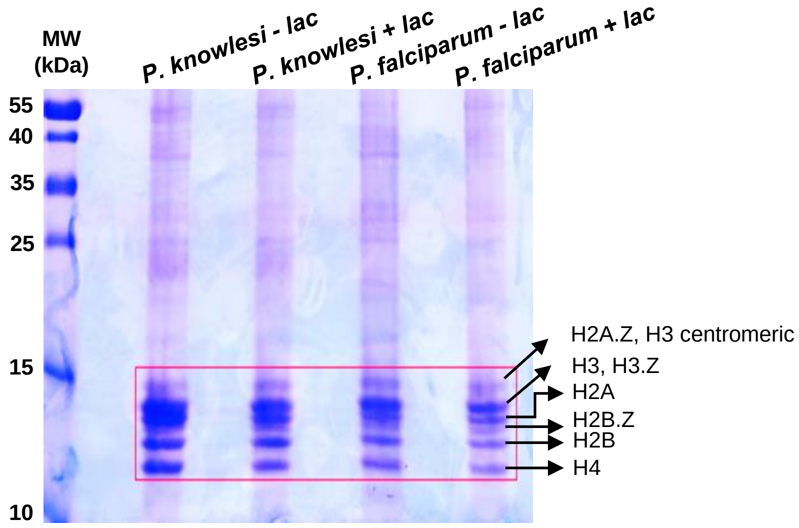


**B**

*P. knowlesi*

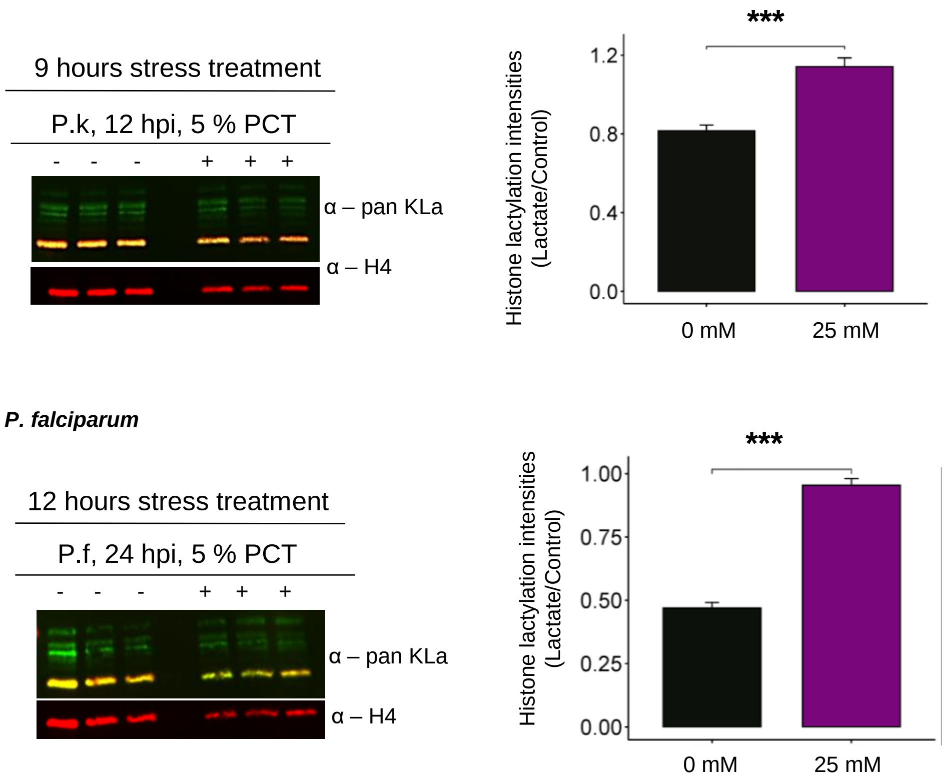

Supplementary figure 5

A

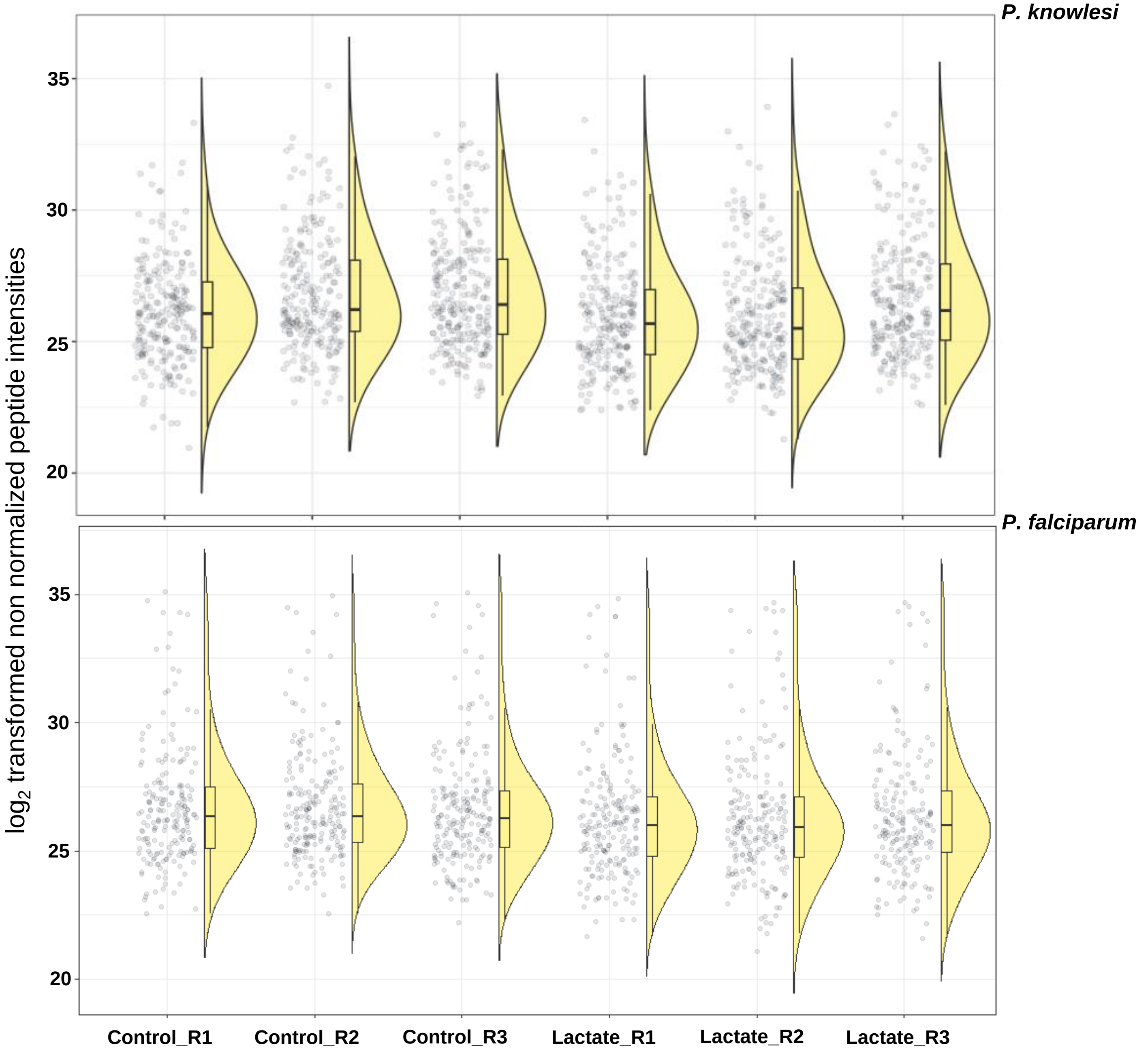


B

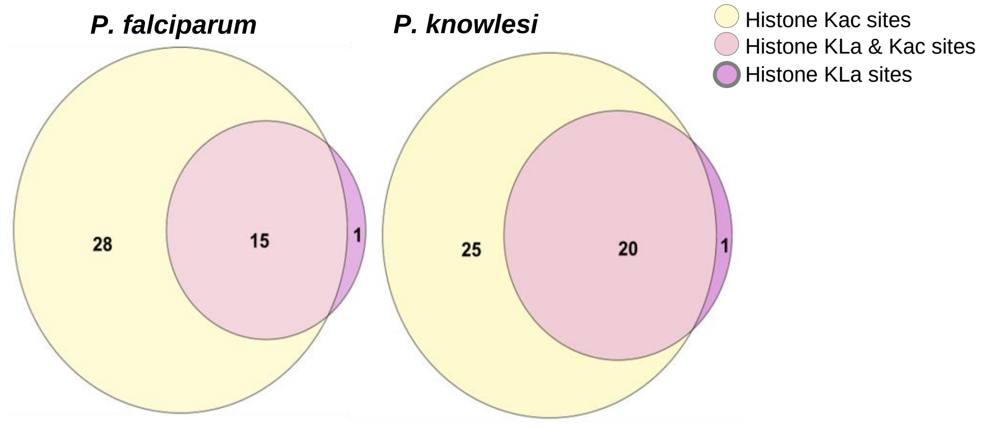

Supplementary figure 6

A

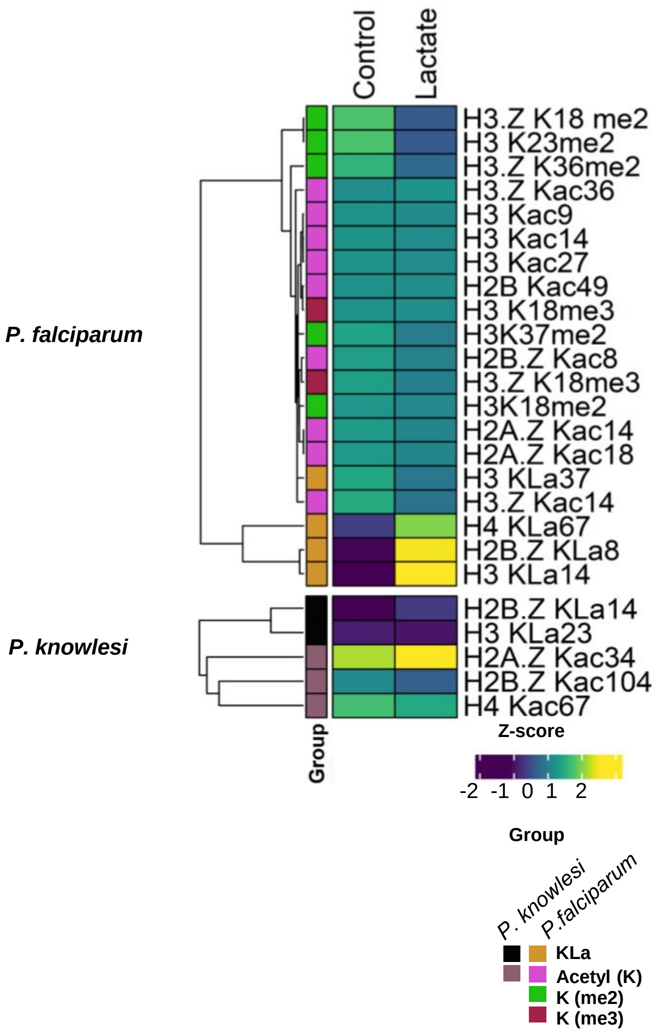


B

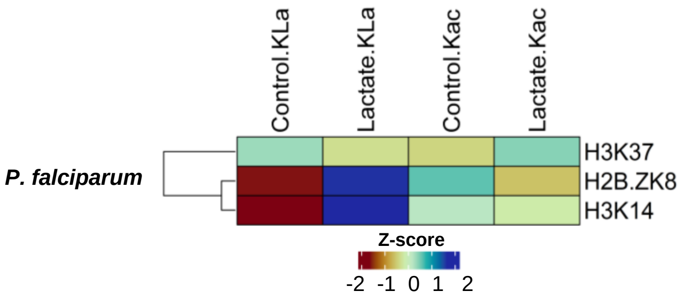

Supplementary figure 7A

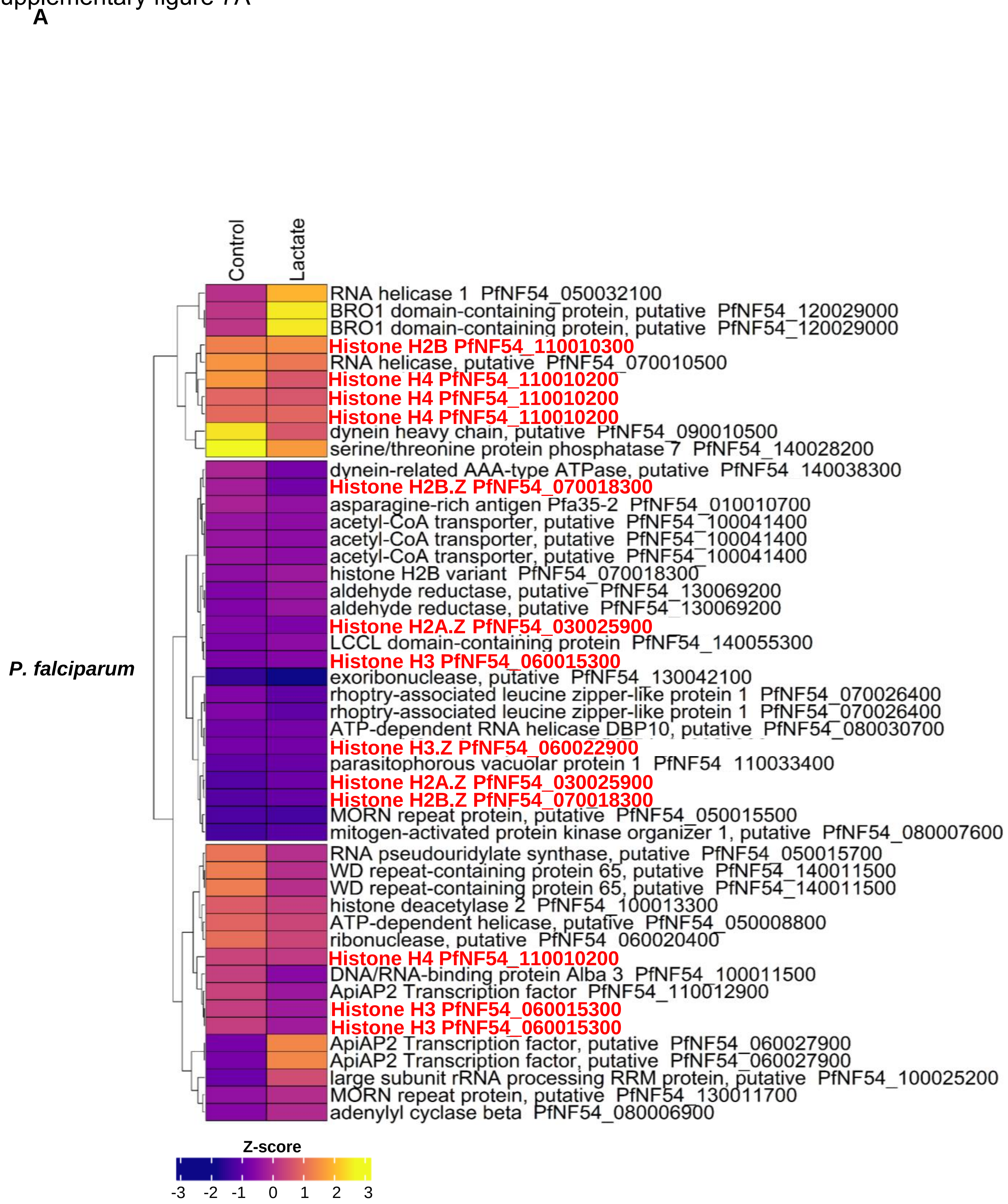

Supplementary figure 7B

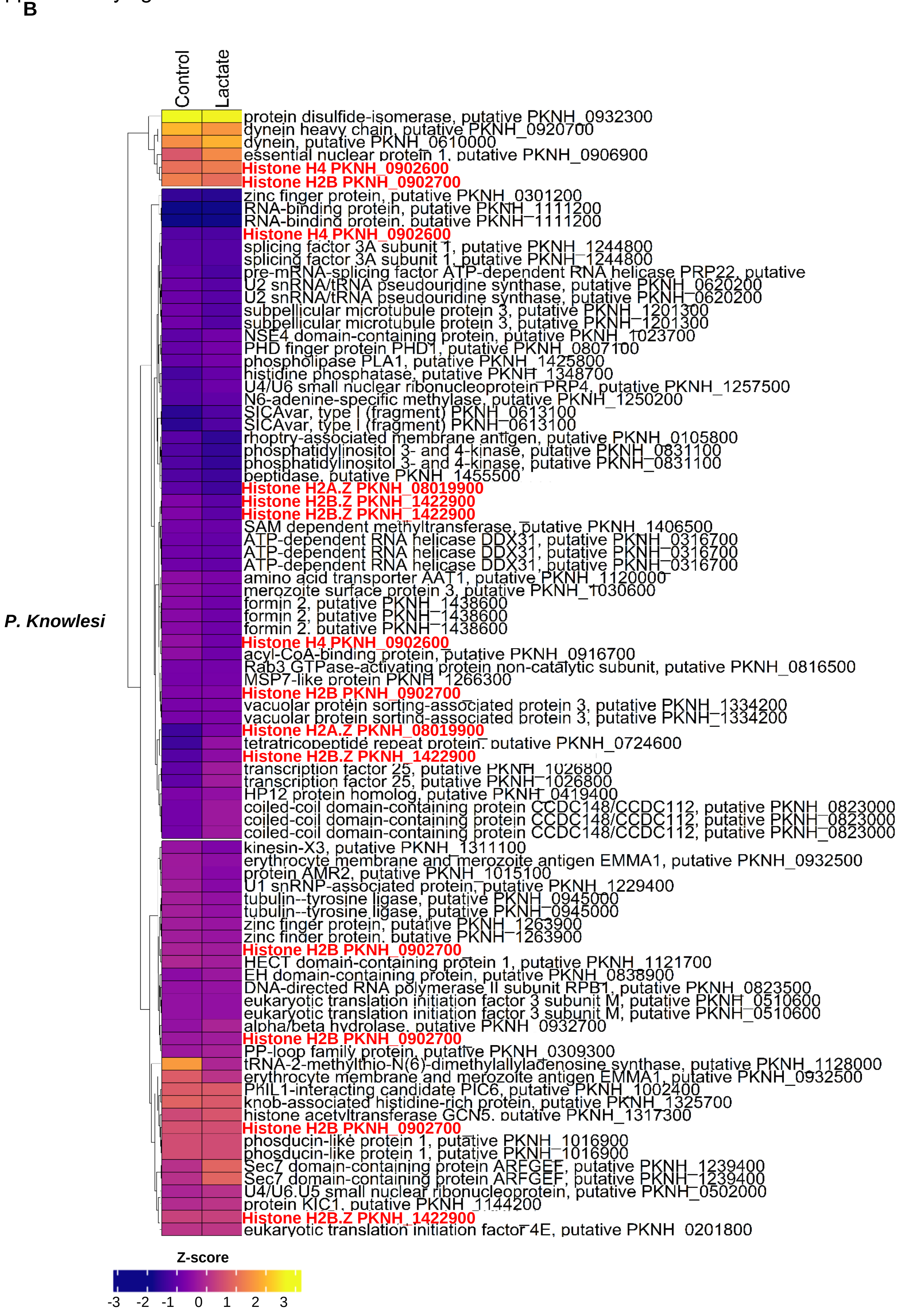

Supplementary figure 8

A

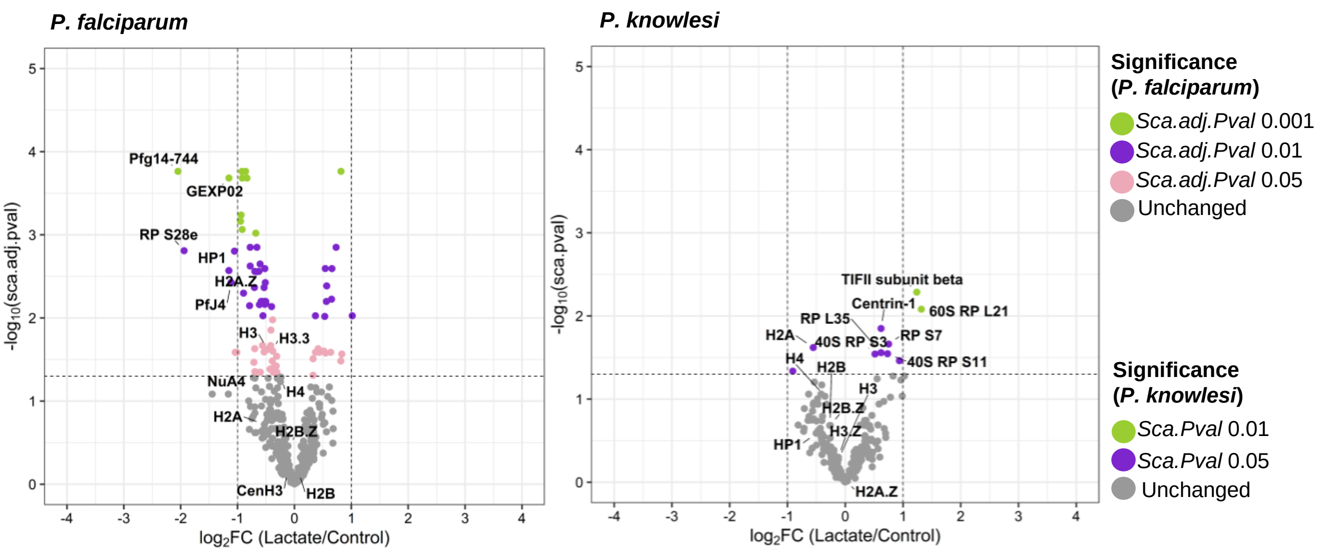


B

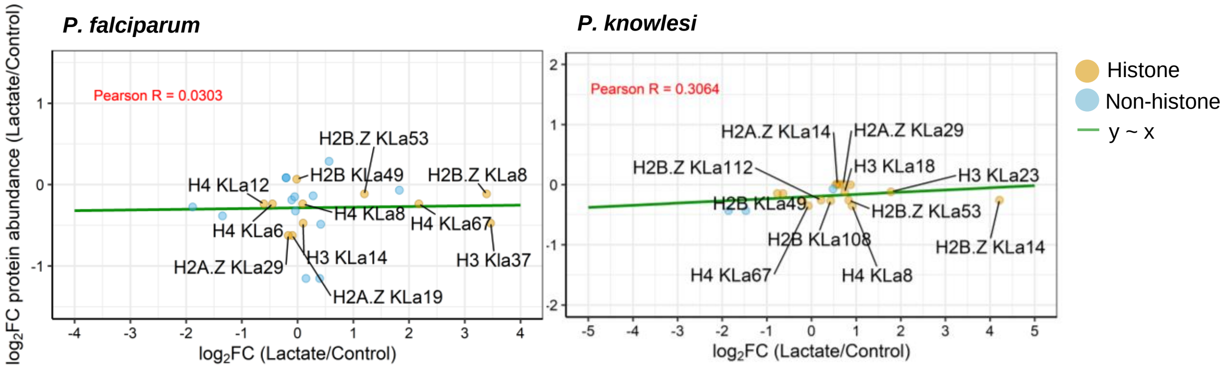

**Biological process**

Supplementary figure 9

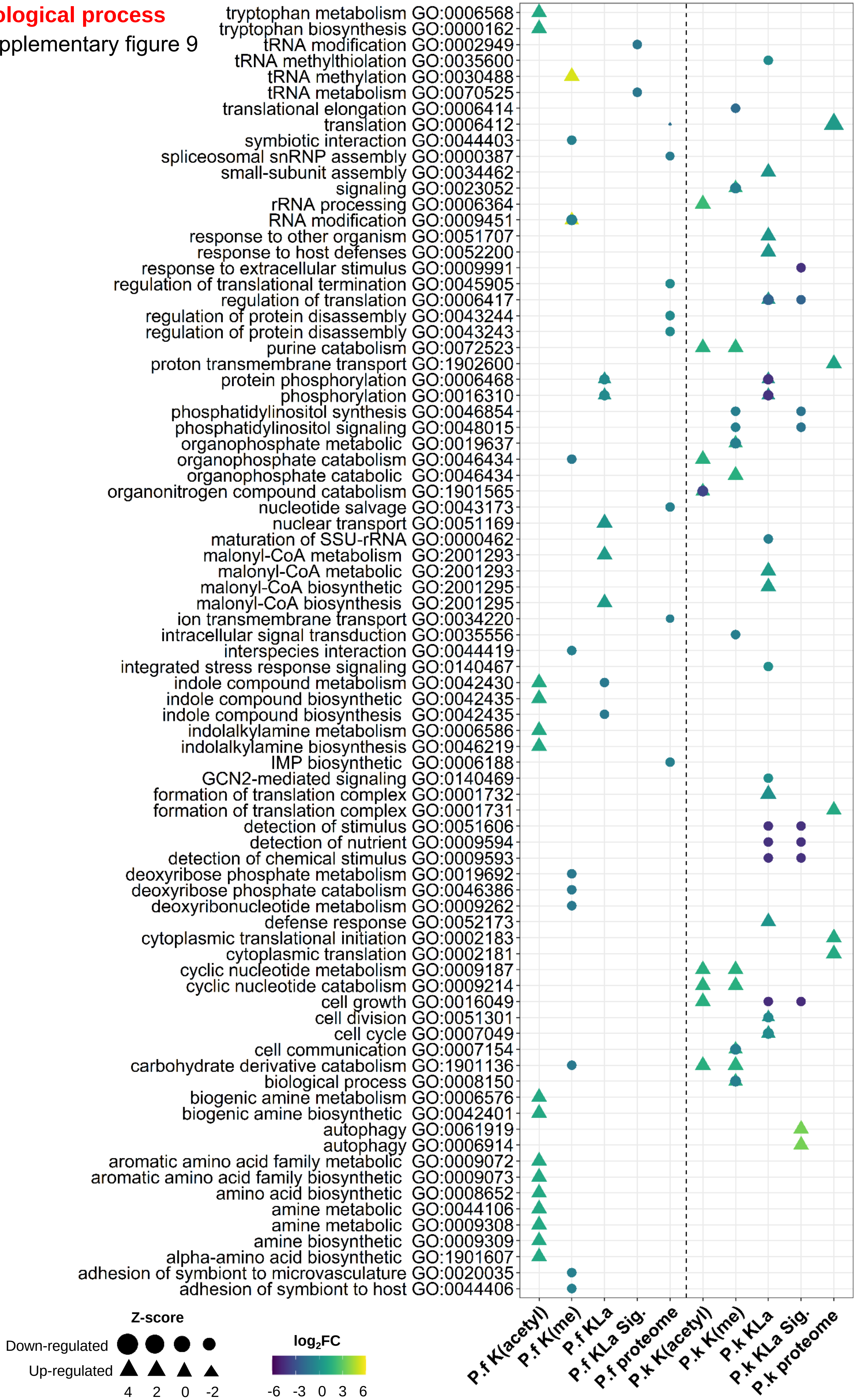

Supplementary figure 10

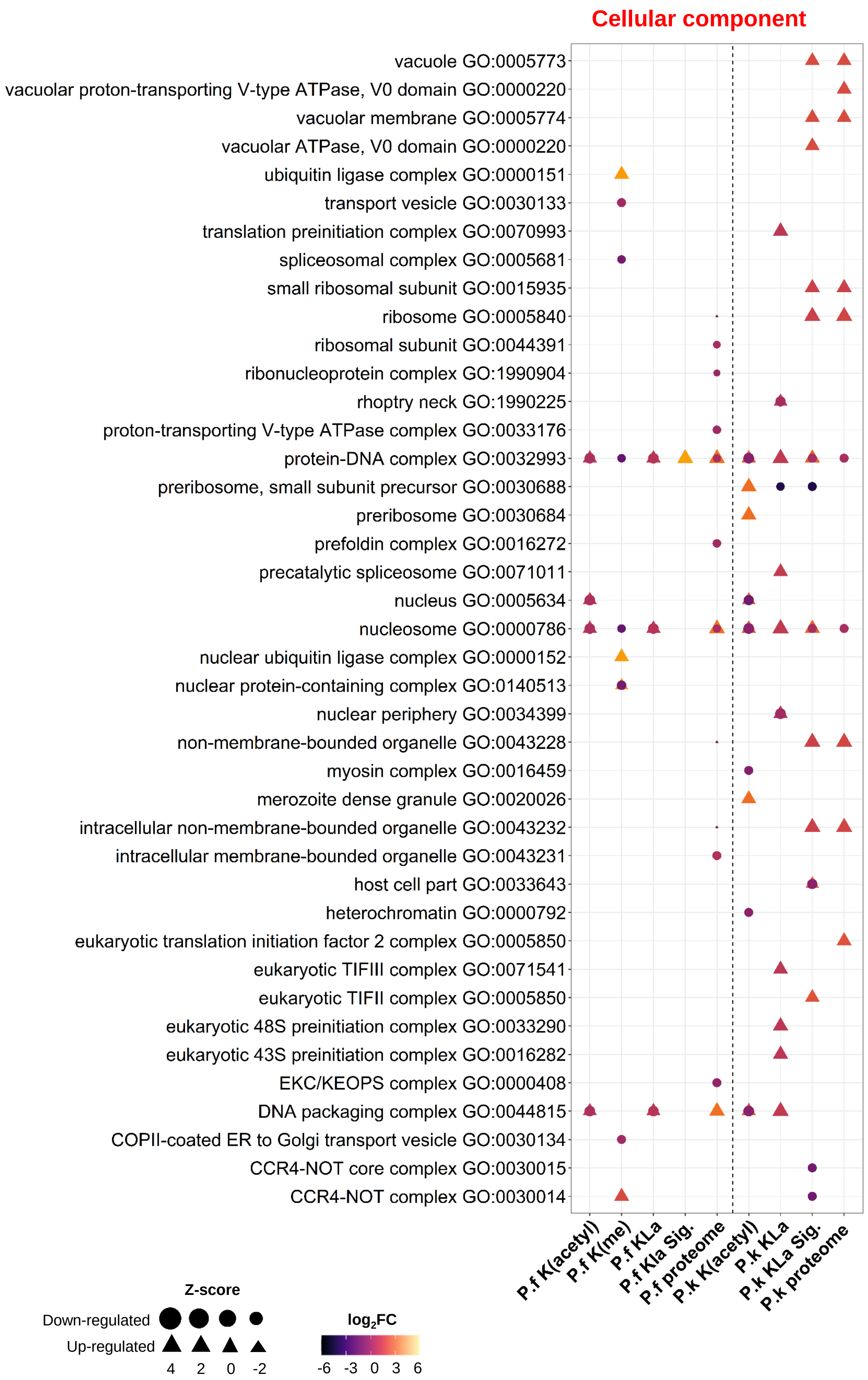

Supplementary figure 11

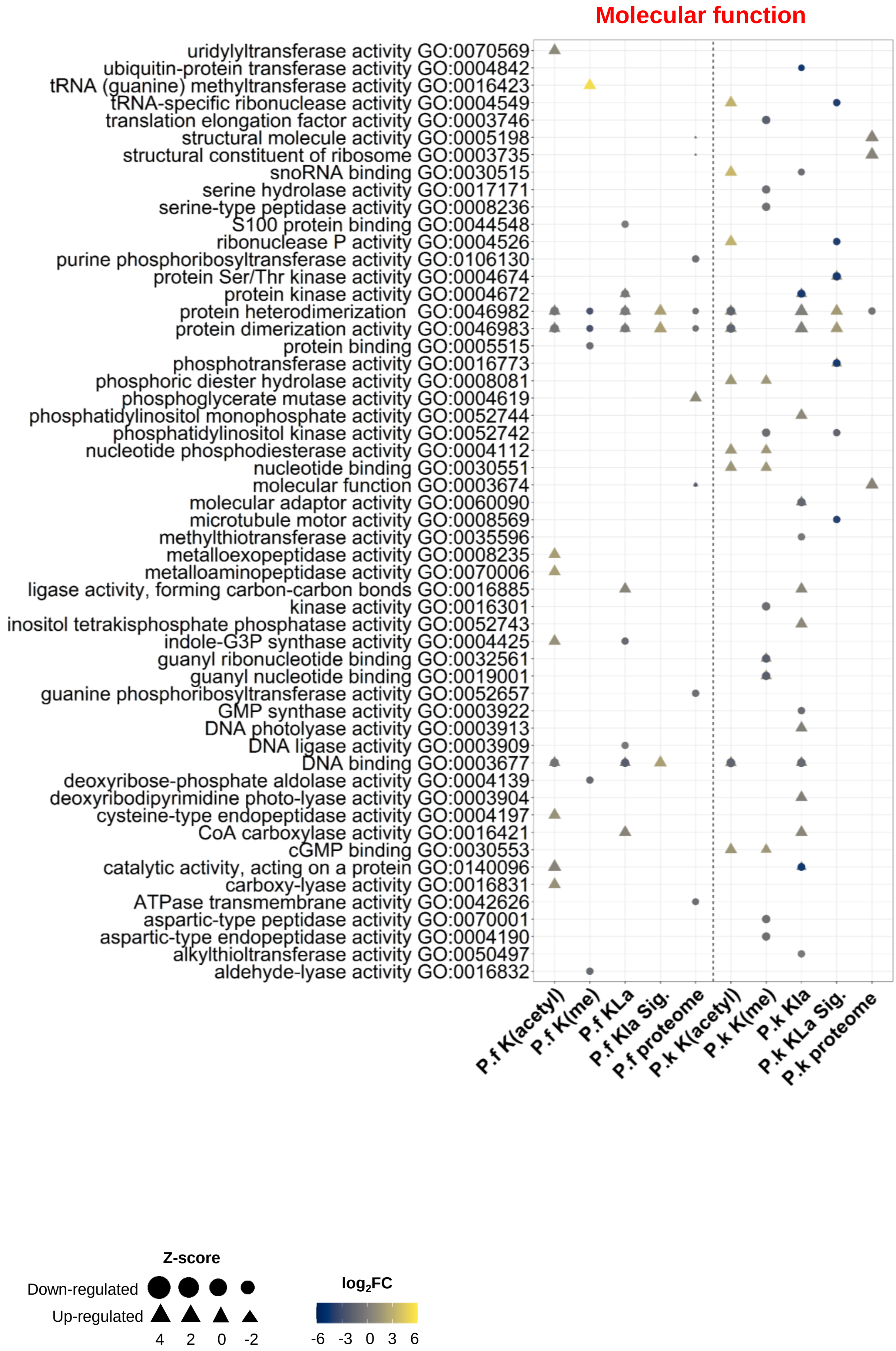

### Supplementary figure 12

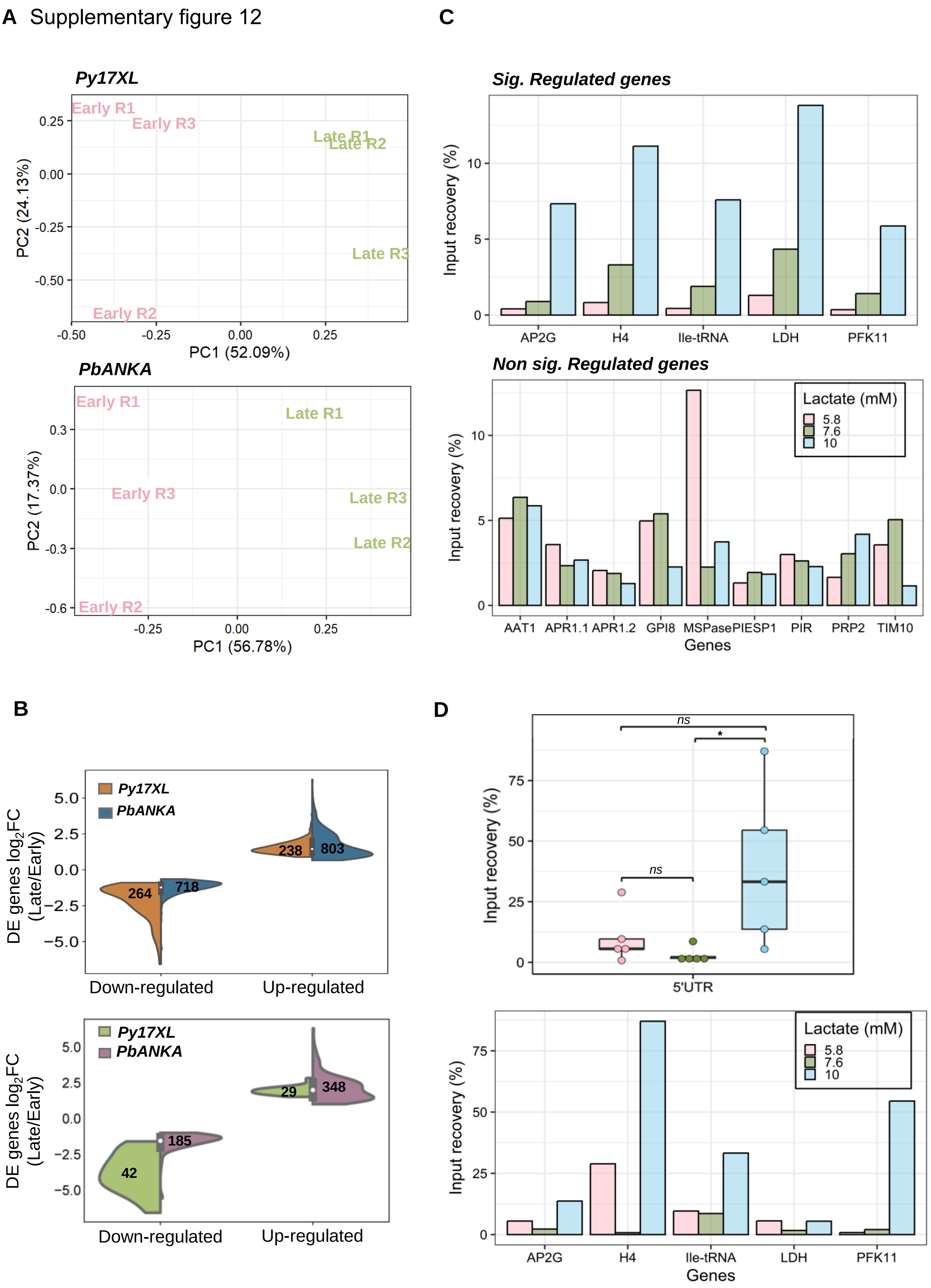

Supplementary figure 13

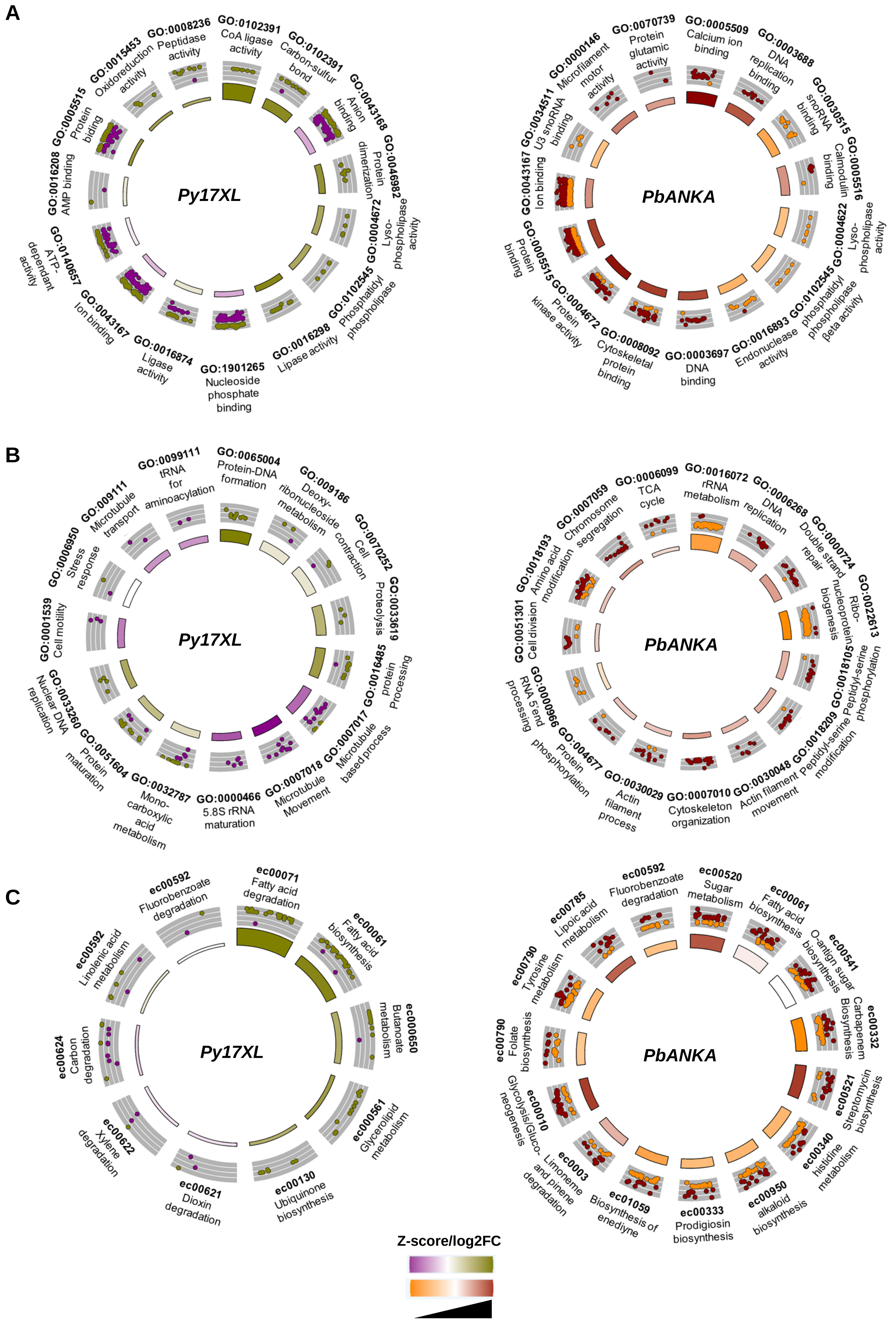
